# Reassessing Deep Learning (and Meta-Learning) Computer Vision as an efficient method to determine taphonomic agency in bone surface modifications

**DOI:** 10.1101/2025.01.31.635872

**Authors:** Manuel Domínguez-Rodrigo, Gabriel Cifuentes-Alcobendas, Marina Vegara-Riquelme, Enrique Baquedano

## Abstract

Recent critiques of the reliability of deep learning (DL) for taphonomic analysis of bone surface modifications (BSM), such as that presented by Courtenay et al. (2024) based on a selection of earlier published studies, have raised concerns about the efficacy of the method. Their critique, however, overlooked fundamental principles regarding the use of small and unbalanced datasets in DL. By reducing the size of the training and validation sets—resulting in a training set only 20% larger than the testing set, and some class validation sets that were under 10 images—these authors may inadvertently have generated underfit models in their attempt to replicate and test the original studies. Moreover, errors in coding during the preprocessing of images have resulted in the development of fundamentally biased models, which fail to effectively evaluate and replicate the reliability of the original studies. In this study, we do not aim to directly refute their critique, but instead use it as an opportunity to reassess the efficiency and resolution of DL in taphonomic research. We revisited the original DL models applied to three targeted datasets, by replicating them as new baseline models for comparison against optimized models designed to address potential biases. Specifically, we accounted for issues stemming from poor-quality image datasets and possible overfitting on validation sets. To ensure the robustness of our findings, we implemented additional methods, including enhanced image data augmentation, k-fold cross-validation of the original training-validation sets, and a few-shot learning approach using model-agnostic meta-learning (MAML). The latter method facilitated the unbiased use of separate training, validation, and testing sets. The results across all approaches were consistent, with comparable—if not almost identical—outcomes to the original baseline models. As a final validation step, we used images of recently generated BSM to act as testing sets with the baseline models. The results also remained virtually invariant. This reinforces the conclusion that the original models were not subject to methodological overfitting and highlights their nuanced efficacy in differentiating BSM. However, it is important to recognize that these models represent pilot studies, constrained by the limitations of the original datasets in terms of image quality and sample size. Future work leveraging larger datasets with higher-quality images has the potential to enhance model generalization, thereby improving the applicability and reliability of deep learning approaches in taphonomic research.

## Introduction

Recently, Courtenay et al. (2024) have cast some doubts on the reliability of some published deep learning (DL) studies used to differentiate bone surface modifications (BSM)^1^, arguing that the high accuracy of these studies are the artificial result of: a) poor quality image dataset, and overfitting of the trained models by blending validation and testing. They replicated the original studies by purportedly using the same model architectures, the same data augmentation procedures, but they modified the training process. In the original studies, the training-validation samples were split at 70%-30%. In Courtenay et al.‘s testing of the original DL models, they divided the sample into training (50%), validation (20%) and testing (30%) sets. Here, we argue that Courtenay et al. (2024) did not replicate any of the three targeted studies, nor did they test generalization shortcomings in the original models. These authors made different methodological decisions about the use of the original data, elaborated different models and extrapolated unjustifiably their results to the original studies, without demonstrating that the original models had problems in generalization. In their logic, there was a disconnection between their *explanans* and *explanandum;* that is, what they set to test and what they eventually proved. Their study, though, is relevant to show that decisions made about methodological procedures have important leverage in analytical outcomes.

The data sets selected by Courtenay et al. were among a few pilot sets used to show the potential of DL in taphonomic analyses. We were aware of their limitations because of the following factors:

a. *Sample size*. DL models inherently require extensive datasets to fully realize their potential and ensure robust performance. However, the datasets compiled for those studies were constrained by an insufficient number of samples in some classes. This limitation arose from the labor-intensive nature of the experimental process, which included experimenting with different agents, cleaning bones, meticulously capturing each bone surface modification (BSM) under the microscope, and subsequently cropping and preparing the images for analysis. For some of the datasets analyzed, this process spanned nearly three years, encompassing project conceptualization, data acquisition, and image preparation. These challenges underscore the practical difficulties associated with generating large-scale datasets for taphonomic research using DL models.
b. *Depth of field*. For these pilot studies, we used a trinocular microscope (Optika SZM-1) paired with a 3 MP digital camera (OptiCam B3). This combo presented problems of depth of field, largely resulting in lack of definition of certain areas of the image. We stressed this problem in the original (Domínguez-Rodrigo et al., 2020), and in subsequent studies (Domínguez-Rodrigo et al., 2024b). This problem affected all BSM and, therefore, it did not bias model performance according to class. This led us to replace this microscope with a Leica Emspira 3 Digital microscope, which is the tool that has provided the bulk of the datasets for our latest studies and provides high-resolution images unaffected by this problem.
c. *Unbalanced datasets*. The first dataset selected by Courtenay et al.-(Abellán et al., 2022)-contained 488 cut marks and 46 crocodile tooth marks. The second data set selected by them-(Domínguez-Rodrigo et al., 2020)-contains 489 cut marks, 103 tooth marks and 63 trampling marks. These datasets are more prone to present generalization issues than more balanced datasets.

It is precisely the awareness of the sample size problem and the highly unbalanced nature of these datasets that led us to consciously make methodological decisions that would boost the performance of the resulting DL models. Our main goal was to develop models that could learn to identify agency. The first of these decisions was not to implement the traditional training-validation-testing split that is the common machine learning protocol (Chollet, 2022). We are aware that separating the training, validation and the testing sets is the adequate analytical process for normal datasets, as we have continuously implemented in our extensive machine learning (ML) experience (Abellán et al., 2022; Domínguez-Rodrigo, 2019; Domínguez-Rodrigo and Baquedano, 2018; Moclán and Domínguez-Rodrigo, 2023). However, having done that triple data split with these small datasets would have presented the following problems:

a. *Reduction of the learning process*. The division of a small dataset into training, validation, and testing subsets significantly limits the amount of data available for training, which is critical for effective generalization in deep learning (DL) models. This is especially so when dealing with unbalanced samples. Effective learning and pattern recognition in DL models require substantial data to adequately capture the underlying features and variability of the dataset. Insufficient training data increases the likelihood of model underfitting, as the model may fail to learn meaningful patterns. Additionally, small testing datasets may result in unreliable performance metrics, as they may not sufficiently represent the data distribution for each class, causing measures such as F1 scores to fluctuate substantially. These limitations also impact the backpropagation process, weight adjustment, and overall learning. The model’s hyperparameters are optimized based on feedback from the validation set, and when the validation set is not representative, the model’s recalibration may be compromised, either due to insufficient learning or overfitting to the validation data. Consequently, in such a situation the model’s generalization performance is also impaired when evaluated on a separate testing set. Therefore, the size and balance of the validation set are as crucial as those of the training set. A poorly-sized or unrepresentative validation set can hinder the model’s ability to learn effectively. Given the constraints of our dataset, which was limited in size and unbalanced, we prioritized larger training and validation sets at the expense of a separate testing set. To mitigate the potential for overfitting and address these challenges, we employed several techniques discussed below.
b. *Increased variance and spurious metrics*. Small subsets can artificially amplify the variance between them. Some subsets may contain outliers or present homogeneous/heterogeneous components, skewing the training or validation process. In such cases, performance metrics become more sensitive and yield inconsistent results across multiple experiments.
c. *Unreliable inferences about the generalization potential of the model*. The use of small and unbalanced datasets compromises the reliability of inferences about the generalization potential of a DL model. An undertrained or underfit model inherently renders testing set metrics unreliable, as it lacks the capacity to learn meaningful patterns effectively. Even if a model is adequately trained, generalization inferences drawn from a testing set can be limited if the set does not manage to be representative enough of the sampled population. With small testing sets, this condition is rarely met. For instance, in Courtenay et al.’s (2024) analysis of Dataset 1, the “crocodile” class testing sample was composed of only 13 images of tooth marks (30% of the sample). Recent research (Domínguez-Rodrigo et al., 2025) has demonstrated the extensive variability in tooth pits alone of this agent, suggesting that such a limited sample cannot adequately represent the full spectrum of the original data, let alone the broader population. Similarly, in Dataset 2, the “trampling mark” class was represented by only 18 images, which is far from sufficient to encapsulate the diverse forms of trampling marks produced by different abrasive agents (Domínguez-Rodrigo et al., 2009). This issue also extends to models that might exhibit strong classification metrics on such limited testing sets. Regardless of whether the metrics are favorable or unfavorable, the small sample sizes prevent a reliable evaluation of the model’s generalization capacity. The underrepresentation of certain classes introduces a significant risk of bias, leaving the broader applicability of the model unverified. This stands in stark contrast to standard DL protocols, which typically involve testing models on hundreds or thousands of images per class, thereby ensuring a more robust and reliable assessment of generalization potential.

By having adopted the triple-split approach, Courtenay et al (2024) may have incurred in deficient modeling (especially for underrepresented classes), since for the first data set, they have trained their DL models, for example, using fewer than 25 crocodile images. A recent study of crocodile tooth pit morphology alone shows that there were a minimum of 64 forms of tooth pits in the same experimental set (Domínguez-Rodrigo et al., 2025). A model loss and backpropagation process is determined by the validation set. In this regard, Courtenay et al. must have used only 6 crocodile images to recalibrate (i.e., update the weights through backpropagation) their models. If using their second dataset, that would have resulted in training the model on 35 trampling marks and validated it with only 9 trampling marks. Needless to say that such reduced validation (and potentially, training) data sets are of little value for DL analysis, since they cannot sample the wide variety of each of those BSM. If all these BSM were from homogeneous and non-overlapping classes, one could agree to operate with such meagre training and validation sets, but given their intense feature overlap, such small sample sizes render the model performance far less reliable than when using larger training/validation datasets, because of deficient training. Such small validation sets propitiate low precision and recall, as Courtenay et al.‘s results show.

In their study, these authors achieved only slightly lower global accuracy on their testing sets than the original studies. For Dataset 1, their accuracy is 92% (in the original study it ranged between 96%-99% according to model [Abellán et al., 2022]). This means that their large cut mark sample was properly trained and classified according to the testing set (91.8 % of precision, perfect recall, and 98.7% of F-1 score). It was only the smaller crocodile data that was unlearnt and misclassified because of the paucity of data for training, validation and testing caused by the triple data split. For Dataset 2, Courtenay et al.‘s replicated testing accuracy was 86% (in the original study it was 90% [Domínguez-Rodrigo et al., 2020]). The larger sample of cut marks was unaffected by the triple data split showing high precision (97.8%), high recall (0.918) and high F-1 score (94.7%), whereas their smaller samples have been learnt more deficiently by the models (tooth marks with a F-1 score of 0.68 versus 80% in the original study), with the smallest trampling subset, being almost completely misclassified (F1-score =36%).

This information alone should reflect the effect of the triple data split on small unbalanced datasets, especially in classes with insufficient sample sizes. This is a confirmation of the “long-tail effect”, in which a DL model performs satisfactorily in classes with more data, but underperforms in classes with substantially fewer data (Rather et al., 2024; Sabha et al., 2024).

If one wants to produce traditional DL models that maximize learning and generalization using small unbalanced datasets, splitting the original dataset in three fixed subsets (with only one for training) is not optimal for model learning. This is where a tradeoff must be selected between learning and generalization. The best way to mitigate the above mentioned issues is either to sacrifice the testing set or to use cross-validation (Brownlee et al., 2021; Hastie et al., 2017; Kuhn and Johnson, n.d.)^2^. The first approach is adequate for model development. In this case, validation can be used as a proxy for testing. The shortcoming of this approach, as mentioned above, is the potential for overfitting on the validation set. To avoid this, data augmentation and regularization methods can be implemented. Transfer learning can also help to leverage existing feature knowledge and fine-tune it on the smaller dataset. This is the approach adopted in the three original studies re-analyzed by Courtenay et al. (2024). It produced fairly accurate models and performance metrics that indicated good precision and F-1 scores for most classes. The main downside is that it cannot be assessed how good the models are at generalizing beyond the validation data sets. Another drawback of this approach is that it is difficult to know if this method produced overfit on the validation set or not. It should not be assumed as a default.

For this reason, the second option (K-fold cross-validation) could be more adequate, and this one was not implemented in the original studies of the three datasets. Cross-validation consists of splitting the original dataset into k subsamples (folds), using one for validating and the others for training. This process is repeated n times, one for each fold created. For example, for a 5-fold cross-validation, the sample uses 80% of the dataset at a time for training and the remaining 20% for validating. By selecting different sets of images for each fold‘s training and validation, the model is exposed every time to different training and validation splits (and image sets). The final result is obtained averaging the accuracy values of each fold. This maximizes data usage while it provides robust validation. Usually, cross-validation is carried out within the training-validation sets and a holdout testing set is used for assessing generalization. Here, we opted to use all data for training-validation through cross-validation, for the same reasons as described above: a representative separate testing set would have substantially reduced the available data for certain underrepresented classes for training. When employing cross-validation for evaluation, the validation set rotates across folds, highlighting potential issues regarding the representativeness of the original data, making a test set a less immediate necessity.

Therefore, when using a small unbalanced dataset, analysts have two options which are very unequal in their results: training-validation-testing or training-validation only (with or without cross-validation). The former reduces available data for training and hyper-parameter calibration through validation, potentially leading to underfit models, higher variance and less stable performance metrics. The latter optimizes the use of data, but may develop overfitting of the validation data, thus not reflecting the true generalization properties of the model. Here, we will test to what extent the original studies were methodologically overfit *versus* to what extent Courtenay et al.‘s remodeling was underfit, by applying this and additional methods to the original datasets.

We will test two hypotheses on the same three datasets used by Courtenay et al. (2024):

*Hypothesis 1*. Image quality has biased the models, probably by different distribution of biasing effects (e.g., brightness, contrast) in different classes. If applying grayscale intensity-augmented methods to the same datasets, this should lead to widely divergent results between the baseline models and the augmented ones.

*Hypothesis 2*. The training-validation split (to the exclusion of a separate testing set) has led to overfitting processes, either on the validation set or on both sets, resulting in artificially high accuracies and low loss. If applying cross-validation methods on the same models and datasets, there should be widely divergent results in the classification metrics of all classes and the overall accuracy of the models. It could be argued that since training of cross-validated models is also carried out on a training-validation split, it has a similar potential to overfit on the validation set as the baseline models. For this reason, implementing a separate and different method (e.g., one-shot or few-shot meta-learning), which can efficiently use separate testing sets on small datasets, should yield widely divergent results if the DL train-validation method is overfit. Failure to show divergent results by this combination of methods implies rejecting the hypothesis. As a final test of this hypothesis, we will also include a new testing set for Datasets 1 and 2 created with BSM that are new and that were not part of the original samples used for the published models. Should the models be overfit, we are expecting a poor performance on the new testing sets.

## Methods and sample

The present study evaluates the reliability of deep learning (DL)-based computer vision (CV) models for classifying experimental taphonomic BSMs. To achieve this, the study utilized three previously published datasets (Abellán et al., 2022; Domínguez-Rodrigo et al., 2020; Pizarro-Monzo et al., 2023), the same ones used by Courtenay et al. (2024), and replicated the analyses originally performed, by implementing additional controlled methods that enabled testing: a) if the quality of images had any impact on model performance, and b) if the training-validation method of the original studies resulted in overfitting through the training and validation feedback. The datasets used in this research include:

1. Dataset 1 (DS1): Published by Abellán, Baquedano, and Domínguez-Rodrigo (2022), available at https://doi.org/10.7910/DVN/9NOD8W (last accessed on 07/08/2023). The dataset is composed of 488 images of cut marks and 45 images of crocodile tooth marks)
2. Dataset 2 (DS2): Published by Domínguez-Rodrigo et al. (2020), available at https://doi.org/10.7910/DVN/62BRBP (last accessed on 07/08/2023). The dataset is composed of 488 images of cut marks, 103 images of tooth scores, and 63 images of trampling marks)
3. Dataset 3 (DS3): Published by Pizarro-Monzo et al. (2023), available at https://doi.org/10.17632/3bm34bp6p4.1 (last accessed on 07/08/2023). The dataset is composed of 629 images of tooth scores, 150 images of cut marks (all new, not recycled from Dataset 2, as wrongly interpreted by Courtenay et al., 2024), and 154 images of trampling marks).

All these datasets have been compiled and put together in a public repository specifically created for this paper: https://doi.org/10.7910/DVN/WUSGSW

### Image quality

The three original data sets used by Courtenay et al were deficient in image quality because of the low-quality of the microscope with which they were taken. Although these authors argued that testing the properties of the images (blurriness, contrast, specularity) is “a fundamental prerequisite in the curation of any dataset”, they fall short of arguing why this has impacted the three datasets that they evaluated. To them, sharp definition of edges, fully focused images, optimal levels of contrast and brightness, and adequate hue-saturation (although these are not applicable to their grayscale images under scrutiny) are mandatory to elaborate reliable “more advanced DL-based analyses”. As a matter of fact, these authorśs adverse reaction to noise led them to apply “an erosional morphological operator […] to clean specular maps” (Courtenay et al., 2024: 391). This is coherent with Courtenay and colleagueś habit of reducing within-sample variance by removing outliers and variance-increasing factors (Courtenay et al., 2021). The downside of this approach of data tweaking is that it frequently under-represents the true variance within the populations from which samples are derived, with the potential of biasing classification credentials of tests and models, and with the only questionable advantage of increasing classification accuracy rates, as they have done in other studies (Courtenay et al., 2021).

If the low quality of the images from the three datasets had impacted the models, the most common consequence would have been a poor model performance. Bad image quality typically decreases the accuracy of deep learning models because it affects the model’s ability to learn features and patterns and, therefore, the ability to generalize from the data. Bad-quality datasets usually impact model performance by failing to detect features; blurry and noisy images usually obscure feature edges and patterns, preventing their recognition. They also impact models by introducing inconsistencies that prevent the model from generalizing well; this is especially relevant when the model learns irrelevant features prompted by noise and distortion. Instead of that, the three datasets used by Courtenay et al. originally produced models with good performance in the original studies (Abellán et al., 2022; Domínguez-Rodrigo et al., 2020; Pizarro-Monzo et al., 2023).

The bad quality of the dataset can only have a positive biasing effect if it primarily affects subsamples from one class over the others, but given that in the three datasets, the images were obtained with the same protocols and instrument, all classes are affected by low quality as shown by Courtenay et al. in the resulting low median values of LoG variance, FFT magnitude and CED for the three datasets. In all classes (depending on dataset) the LoG values fall <50% (except one). Courtenay et al argue that cut marks “emerge as the category with the highest amount of images out of focus”, but this results from the fact that the cut mark subsample is the most abundant within the unbalanced original datasets. It has the highest amount of bad-quality images, but also the highest amount of better-quality images, depending on the dataset.

In contrast, although we concur with Courtenay et al. that better-quality images would have been desirable, and have done so by using a better microscope in our recent datasets, we see the advantage of having used these exploratory limited datasets, because the alternative explanation to the good-performing models obtained in the original studies may probably lie in having exploited the advantages of blurry, out-of-focus, insufficient levels of contrast, and saturated brightness of portions of the images as a regulatory method. This is actually what is done when applying data augmentation. Data augmentation consists of adding synthetic noise to the images by blurring them, cropping them, zooming-in-out of them, and distorting their contrast and brightness. This prompts the training model to focus on the features that are relevant to each class, which in most images of the three original datasets appear well defined in the center of each image. Given that we had also applied shear augmentation, the location of the focused area also varied among images, which increased the sample variability and limited potential degrees of differential focusing with specific classes. This, added to the fact that CNNs are known to be invariant to the location of the features within the images (Goodfellow et al., 2016; Rosebrock, 2017; Chollet, 2022) limits the likeliness of the CNN models using the location of the focused areas of the images to make a prediction, rather than the actual features found within those areas. Moreover, data augmentation leads the model to learn in a more robust way from poor-quality images. If this is done in combination with transfer learning, this contributes to learning patterns even better. Courtenay et al.‘s Grad-CAM images only display what their models perceive on the selected images, not what the original models identified, making their insights only applicable to the models produced by them. Here, we argue that the different training strategies followed by Courtenay et al in comparison to the original studies could explain the GRAD-CAM behavior reported in their work. Likewise, the Grad-CAM method is not adequate when dealing with BSM, whose features are not detached or clearly delineated from its contextual surroundings. See an extended discussion of this in the Appendix.

Also importantly, we do not believe that the image-quality analyses presented by Courtenay et al. are necessarily a good representation of the adequacy of the microscopic images used in the original studies. We believe that microscope images cannot directly adapt to standards and thresholds commonly associated with natural non-microscopic images. In this sense, there is a difference between optimal values for natural images, and optimal representation of taphonomic features. One of the most evident cases in which these differences manifest is the utility of specularity when imaging relevant taphonomic features. For example, proper detection of internal microstriations (crucial to identify cut and trampling marks), may require that such inconspicuous features are emphasized thanks to the specular highlights that they produce when optically documenting the BSM. The benefit of highlighting such shallow features can be better appreciated when considering trampling marks, where internal microstriations can be so shallow that they may not be visible in the final image if not for the specular highlights they produce. Moreover, specular reflections can also highlight the outline and trajectory of these BSM. In this sense, the specular highlights are not breaking useful patterns like edges and textures, but they are enhancing how some of these features are documented and allowing the models to use features that would have otherwise not been recorded. From this point of view, adequate representation of taphonomic features in BSM images may conflict with common standards associated with photographically optimal images (i.e., with optimal levels of contrast and brightness). As long as the discriminant features are properly highlighted, other parts of the image may remain out of focus, blurred and poorly defined, thus not affecting feature detection by the CNN models. Given that Courtenay et al have not shown that image quality had a biasing effect in the performance of the three datasets that they tried to replicate, here we will test both interpretations in the form of Hypothesis 1 (outlined above). We hypothesize that if image quality biased the original models, a re-analysis using controlled noise methods consisting of even distortion of brightness, contrast, and sharpness across classes should yield significantly lower accuracy rates and worse performance metrics than the original models published by Abellán et al., (2022), Domínguez-Rodrigo et al. (2020), and Pizarro-Monzo et al. (2023). For this purpose, we will use grayscale intensity-augmentation methods in addition to image normalization.

We generated a function that randomly adjusts the brightness, contrast and sharpness of the image to introduce various variations, that improve the robustness of the model to changes in image lighting conditions.The process implements a data augmentation pipeline tailored for three-channel grayscale images, designed to enhance variability and improve model robustness during training. The function applies random intensity-based—brightness and contrast— and sharpness augmentations independently to each channel, despite the input image being grayscale. By doing so, we make sure that the random augmentations to each of these image features are not too extreme, which could make them too dissimilar from the overall sample, hindering the model’s ability to learn generalizable patterns. For brightness adjustments, pixel intensities in the selected channel are scaled by a random factor between 0.8 and 1.2, simulating variations in lighting conditions. Contrast augmentation modifies the difference between pixel intensities relative to the channel’s mean intensity, with a scaling factor also randomly chosen within the same range, thereby mimicking diverse imaging conditions.

Sharpening is performed using a convolution operation with a predefined kernel, amplifying edges and enhancing fine details to simulate sharper imaging scenarios. Each of these augmentations operates within the valid pixel range (0–255), ensuring no overflow or underflow. The augmented image is then converted to a float32 data type and preprocessed using ResNet50’s “preprocess_input” function for Datasets 1 and 3, which normalizes the pixel values according to the statistical distribution expected by the model (e.g., mean-centered and scaled based on the ImageNet dataset). For Dataset 2, the augmented image is then converted to a float32 data type and preprocessed using VGG16’s “preprocess_input” function, which normalizes the pixel values by subtracting the channel-wise mean values derived from the ImageNet dataset, ensuring compatibility with the statistical distribution expected by the model. By introducing random, channel-specific augmentations to grayscale images while maintaining a three-channel structure, this approach enables compatibility with pre-trained models like ResNet50 of VGG16, improving generalization and robustness in downstream deep learning tasks.

The original dataset used grayscale images on the three channels, instead of a single channel. The reasons to do so was model compatibility. Since transfer learning was used and the original pre-trained models were designed for three-channel inputs, this channel structure was kept for adjustment. Pre-trained weights can be used effectively this way because the input shape matches the expected format. This is an advantage when using grayscale intensity augmentation, because the artificially-induced distortions to brightness, contrast, and sharpness can be independently applied to each channel bringing a much wider range of unique combinations to each altered grayscale image than if we were using a single channel (Chatfield et al., 2014).

### Model architecture

In the present work, we are not targeting the selection of the best performing model, and we are not using ensemble learning. For this reason, we will use single models with each dataset to test the two hypotheses outlined above. Given the overall good performance of ResNet50 in some of our previous works (Abellán et al., 2022; Domínguez-Rodrigo et al., 2024b, 2020; Pizarro-Monzo et al., 2023), we will use this architecture here for Datasets 1 and 3. The reason for using it with Dataset 1 is that in the original study it also provided the lowest loss of all the transfer learning (TL) models (Abellán et al., 2022). The reason for using it with Dataset 3 is that it was the model with the highest accuracy. For Dataset 2, VGG16 provided the best accuracy and loss scores of all the transfer models tested (Domínguez-Rodrigo et al., 2020). For this reason, we will use the same model here.

For the present re-analysis, we will use TL only. The base model consists of the original ResNet 50 (or VGG16) model excluding the top (fully connected and classification) layers and retaining only the convolutional base for feature extraction, as it is the common protocol for TL use (Brownlee, 2017). The original models were trained on the ImageNet dataset, which enables it to be successful under domain variation and adaptable as a general-purpose feature extractor. All the layers of the model were frozen to ensure that the pre-trained weights were not updated during the training of the new domain dataset. The additional following custom layers were added: a Flatten layer converting the feature map output by the TL models into a 1D vector for use with a Dense layer; a Fully Connected Layer with 128 neurons, ReLu activation and He uniform initialization to enable task-specific feature-learning; a Dropout layer as a regularization method to avoid overfitting, dropping 30% of the nodes during training, and a Dense output layer with a softmax activation function. For Dataset1, the ResNet 50 model was compiled with Mini-batch Stochastic Gradient Descent (SGD) as optimizer (learning rate= 0.001; momentum= 0.9), categorical cross-entropy as the loss function, softmax as the activation function (to obtain class probabilities), and accuracy as the metric to evaluate the modeĺs performance during the training phase. For Dataset3, we modified the optimizer from SGD to Adagrad, because we realized that the authors of that dataset had not explored the improvements brought by this optimizer over SGD for their particular image set (Pizarro-Monzo et al., 2023). The VGG16 model was modified by including the use of the “swish” activation function and also the Adagrad optimizer, since we observed in other experiments that the VGG models performed better in general when using this combination (Domínguez-Rodrigo et al., 2021). Given the multiple-class nature of the dataset where VGG16 was used, we used categorical cross-entropy as the loss function and accuracy as the evaluating metric for training.

Prior to DL analysis, the original image datasets were split into training (70%) and validation (30%) sets. In the present work, we did not use hold-out testing sets because of the very small size of the dataset (see discussion above). Models were trained with images of 400 x 80 pixels and data augmentation. Augmentation procedures included random shifting (0.2), shear (0.2), zoom (0.2), horizontal flipping and rotation (40°). The image pre-processing function was the original from the ResNet 50 (or VGG16) transfer learning models. The preprocessing function (commonly known as “preprocess_input” from the Keras “tensorflow.keras.applications” module) includes color normalization for color images, depending on which TL model is used. For some models, this function performs two main operations on the input images: a) The first operation involves scaling the pixel values to a specific range, converting them from the original [0, 255] range to the range required by the CNN ([-1, 1] or [0,1]); b) The second operation normalizes the color channels by subtracting the mean RGB values computed from the ImageNet dataset and dividing each channel by its corresponding standard deviation, ensuring the pixel values align with the statistical distribution expected by each model. For VGG16, the only operation also adjusts the color channels by subtracting the mean RGB values computed from the ImageNet dataset ([123.68, 116.779, 103.939] for R, G, and B channels, respectively) without dividing by the standard deviation, ensuring the pixel values align with the input expectations of VGG16. For Resnet 50, this operation is done also subtracting the mean RGB values computed from the ImageNet dataset for the R, G, and B channels (Keras 2 API documentation [https://keras.io/api/applications/resnet] accessed on January 23, 2025). Therefore, the pre-processing function is also taking image normalization into account. When using grayscale intensity-augmentation, this preprocessing functions were also embedded in the augmentation function.

The main novelty between the original models and those re-trained here as baseline models consists of the use of regularization methods (i.e., Dropout). Training was performed in batches, which varied according to the original models to adapt as closely as possible to the original configuration of the published models. For Dataset 1, training used batches of 64 images and validation used batches of 32. For Datasets 2 and 3, training used batches of 32 images and validation used batches of 32, since that is how they were originally coded.

The model fitting involved training during 100 epochs. CNN models were developed using the Keras API with a Tensorflow backend, and computation was performed on a Nvidia Quadro P5000 GPU(HP Z6 Workstation) within a CUDA computing environment.

### Analytical process and comparative framework

A five-stage process was implemented:

1. The first stage was to determine model performance using a training-validation set to generate a baseline model.
2. The second stage enabled contrasting hypothesis 1 (image quality impacted the baseline model performance). Here, the image datasets were transformed through grayscale intensity-augmentation by introducing distortion in image sharpening, brightness and contrast for all classes equally. New models were generated to compare with the baseline models.
3. The third stage consisted of determining if the sample splitting method overfit the models. For this stage stratified cross-validation methods were applied, given the unbalanced nature of the datasets.
4. Given the small sample size, to exclude the potential of validation overfitting in the previous modeling, an independent testing set was used through a different deep-learning method involving model-agnostic meta-learning (MAML), which is part of the few-shot learning array of methods. These methods enable learning through the use of small datasets and, therefore, the possibility of testing models with independent testing sets, regardless of sample size.
5. For Datasets 1 and 2, a new set of BSM was used as a separate testing set. This separate set is composed of recent images that were not part of the original study. This is an ultimate test to the validity of the models and the lack of methodological biases. This was not applied to Dataset 3 because: a) we did not have additional BSM to fulfill 30% of such an extensive dataset; b) all our carnivore tooth mark samples had been used in that study; c) the original study was large enough to generate a testing set, and d) also because its classification metrics derived therefrom were not affected by potential methodological biases, as shown in Courtenay et al.‘s (2024) re-analysis. The reliability of the modeling for this data set (and the other two) is ultimately addressed by using meta-learning, with independent testing sets.

### K-fold cross-validation

A stratified k-fold cross-validation was implemented to make sure the different classes in the unbalanced samples were proportionally represented in each split/fold. The “StratifiedKFold” function from scikit-learn library was used to ensure a 75% (training)-25% (validation) split rotated through all the images after 4 folds. The function generates unique splits for each fold. In each fold, the unique indices generated for training and validation are mutually exclusive, resulting in unique images for each set. Over all folds, every image will appear only once in the validation set and the remaining times in the training set. This results in every image used for validation only once across k folds. Averaged cross-validated performance metrics were obtained at the end. To test the impact of the training-validation method over the baseline model, this was applied without grayscale intensity-augmentation for comparative purposes.

### Model-Agnostic Meta Learning (MAML) method

If the baseline models had been overfitted during training, it would be expected that a separate testing dataset using a different method would show this by providing lower accuracy and higher loss than the trained DL models, as well as low values for the performance metrics (precision, recall, F-1 score). To test Hypothesis 2, in addition to the cross-validation method above, we implemented a few-shot analysis, which enables the use of small datasets for learning and the training-validation-testing splits that are the protocol of machine learning methods-and more specifically, deep learning methods using larger samples. We implement this approach here, with a triple data split method, as Courtenay et al., instead of a dual training-validation set. The difference is that we are doing it through a model/method that has maximized its learning, instead of restricting it, compared to the baseline models used by Courtenay et al.

There exists a diverse range of few-shot learning (FSL) methods, many of which differ significantly in philosophy and structure. These include siamese networks, prototypical networks, relation networks, matching networks, and model-agnostic methods (Jadon and Garg, 2020; Ravichandiran, 2018). Among these, Model-Agnostic Meta-Learning (MAML) has been selected as the method of choice for this study due to its independence from specific model architectures. MAML’s design enables it to adapt to a wide variety of model structures without substantial modifications (Finn et al., 2017; Liu et al., 2024; Ravichandiran, 2018). The core of MAML involves a meta-learning process in which model parameters are fine-tuned and updated, rendering them highly adaptable to novel models and tasks.

In MAML, the concept of “task” forms the foundation of its meta-learning approach. Tasks represent individual learning problems or datasets that the model must address. In the context of the FSL method employed here, each task comprises a small dataset with limited examples, where the model learns from these constrained data. Tasks may involve various classification problems, such as classifying images into different categories, each using a variable number of subsamples and classes. Across tasks, the number of examples per class can also vary.

MAML’s meta-learning process operates over multiple tasks, training the model to quickly adapt to any new task it encounters. This approach contrasts with the meta-learning strategy employed in deep convolutional neural networks (DCNNs).

The MAML methodology operates through a two-stage learning process. The first stage involves task-specific adaptation, where the model’s parameters are updated using gradient descent based on the data from a specific task. During this stage, the model begins with a shared set of parameters, which are fine-tuned using one or more gradient descent steps applied to the task’s support set (training data). In the second stage, task-specific parameters from multiple tasks are leveraged to further adjust the shared parameters using a query (evaluation) set. This iterative process optimizes the initial parameters, enabling the model to generalize effectively across tasks. The ultimate goal of this process is to learn an initialization that supports efficient adaptation to new tasks with minimal additional training data. As the number of tasks increases, the parameters are refined to maximize performance. In essence, the first stage focuses on minimizing task-specific loss through optimization, while the second stage refines the parameters through meta-learning to enhance generalization across unseen tasks.

In this study, MAML was implemented within the framework of transfer learning (TL). Here, the ResNet50 model was employed as the base model for MAML. To retain the pre-trained features, all layers of these models were frozen. The MAML model was constructed with a sequential architecture, incorporating the TL models and additional overlaid layers. These layers included a convolutional layer with 512 filters and a kernel size of (3 x 3), a Global Average Pooling layer to reduce feature map dimensions, a fully connected dense layer with 512 units, a Dropout layer, a Batch Normalization layer, and a final output dense layer with Softmax activation. Regularization strategies were employed to mitigate overfitting, including a 60% dropout rate (twice as high as the earlier DL models) and early stopping, monitored through validation loss with a patience threshold of 15 epochs and weight restoration.

Image preprocessing included normalization using the preprocessing functions of the TL models. The dataset was partitioned into training (70%), validation (15%), and testing (15%) subsets, resulting in 374 training images and 160 images evenly split between validation and testing for Dataset 1. For Dataset 2, training involved 458 images and validation-testing used 193 images, split between the two. Dataset 3 was composed of 653 images for training and 280 images split into two validation-testing sets. Data augmentation techniques such as random shifting, shearing, zooming, horizontal flipping, and rotation were applied, in addition to image standardization to 400 x 80 pixels.

The MAML model used the Adam optimizer with a learning rate of 1e-03. Loss was measured using “sparse categorical cross-entropy” since labels were one hot encoded. Training was conducted over 100 epochs with a validation interval of 1.

The study employed variable task and shot configurations, categorized as low (few shots and more tasks), and high (more shots and few tasks). For the Dataset 1, the high-shot configuration involved 374 images (10 images x 2 classes x 19 tasks), which remained at about the total number of training images. No low-shot task was implemented because of the positive results of the high-shot approach. For Dataset 2, the high shot-task configuration was based on 458 training images (10 shots x 3 classes x 15 tasks). The low-shot configuration was 5 shots-30 tasks. For Dataset 3, the high shot-task configuration was based on 653 training images (10 shots x 3 classes x 20 tasks). The low-shot configuration was 5 shots-40 tasks.

To ensure stability, the task-shot module was configured with “replace=False,” preventing resampling. The restricted configuration employed here produced more stable models with reduced or no overfitting, depending on the set. The two configurations (low, high) facilitated a comparative analysis of accuracy improvements driven by varying task-shot combinations.

### Testing the original models with novel testing datasets

To the traditional analyst, nothing certifies better the validity of a model than testing it against a sample of unseen data (i.e., testing data not used for training or validation) (Chollet, 2022). For taphonomy, given that such extensive data require additional time-consuming experimentation, we borrowed a new set of experimental BSM from recent (unpublished) experiments. For Dataset 1, we managed to get access to a recent small collection of crocodile tooth marks. The original sample had been obtained with crocodiles in Faunia (Madrid) (Baquedano et al., 2012; Domínguez-Rodrigo and Baquedano, 2018). The new testing set was obtained from an experiment conducted by our colleague Edgard Camarós at Altamira Zoo (Santander, Spain). Two adult dwarf crocodiles (*Osteolaemus tetraspis*), both male, were utilized in this experiment. Carcasses, comprising partly defleshed limbs of adult pigs and a pelvis, were collected after 10 minutes of feeding exposure. The bulk of the tooth marks obtained were tooth pits, which were not used in the present study; however, a small number of tooth scores were photographed with an Olympus LEXT OLS3000 confocal microscope and will be used here. This was complemented with a few tooth scores of the Faunia collection that had been discarded in the original study because of the poor resolution of the images obtained with the Optika microscope. Here, they were documented with a Leica Emspira 3 optical microscope. A total of 20 new tooth scores were gathered using both sources. This number equals 44% of the original complete sample used by Abellan et al. (2022), and is slightly bigger than Courtenay et al.ś (2024) original testing sample of the same dataset (n=13).

We also incorporated a set of 65 cut marks from a recent experiment (Cifuentes-Alcobendas, 2025), different from the original dataset used by Abellán et al. (2022). This new sample was photographed also using a Leica Emspira 3 microscope. We decided to keep the cut mark sample proportionally smaller than the original study, to better assess the potential biasing effect of the crocodile tooth mark under-represented class, purportedly most affected by analytical biases.

For Dataset 2, we used an additional set of cut marks from Cifuentes-Alcobendas (2025), given that the cut mark sample for both original studies was the same (Domínguez-Rodrigo et al., 2020; Abellan et al., 2022). Again, given that the cut mark samples for both datasets originally scored high in the classification metrics, we wanted to minimize their effects on the more biased underrepresented classes. It is for this reason that we kept, therefore, a small cut mark testing set. Additionally, we included 25 new trampling marks randomly selected from the enlarged sample generated by Cifuentes-Alcobendas (2025). This would equal about 40% of the original trampling sample used for dataset 2 (n=63) (Domínguez-Rodrigo et al., 2020). For the tooth mark sample, we used a testing set of 30 tooth marks from wolves, which were never included in the original study of Dataset 2, and which were acquired through microscopy using a Leica Emspira 3. This equals almost 30% of the original tooth mark sample used for the analysis of Dataset 2 (Domínguez-Rodrigo et al., 2020). We were careful not to introduce tooth marks from different carnivores from which the original model had been trained (lions and wolves), given that they are different enough to be classified by class and can be potentially misclassified (Domínguez-Rodrigo et al., 2024b; Jiménez-García et al., 2020).

For Dataset 3, we did not implement any additional testing sample for lack of new BSM, which could be used in the proportion required as a testing set for the larger original sample used for that BSM assemblage (Pizarro-Monzo et al., 2023). However, it was also not necessary, since the replication carried out by Courtenay et al. (2024) with an additional testing sample split already showed good classification (except for the trampling subset).

## Results

### Dataset 1

A trained ResNet 50 model yielded an accuracy on the classification of the validation set of 98.75% (loss= 0.0251). High accuracy and low loss together suggest that the model generalizes well and is not overfitting or underfitting. The results show that the model is both accurate and calibrated in its predictions. The performance metrics indicate high precision, recall and F1-score for both classes (Table 1). Only 2 BSM (2 crocodile tooth marks) out of 160 were misclassified from the validation set (n=160) (Table 5). Up to this point, the influence of potential overfitting on the validation set cannot be ruled out.

**Table 1.** Classification report of the performance metrics from Dataset1 (support set= 160). Compare these results with those of Courtenay et al. (2024: Table 3).

**Baseline model**
|  | precision | recall | f1-score | support |
| --- | --- | --- | --- | --- |
| crocodile | 1 | 0.86 | 0.92 | 14 |
| cut | 0.99 | 1 | 0.99 | 146 |
| accuracy |  |  | 0.99 | 160 |
| macro avg | 0.99 | 0.93 | 0.96 | 160 |
| weighted avg | 0.99 | 0.99 | 0.98 | 160 |

**With grayscale intensity augmentation**
|  | precision | recall | f1-score | support |
| --- | --- | --- | --- | --- |
| crocodile | 1 | 0.86 | 0.92 | 14 |
| cut | 0.99 | 1 | 0.99 | 146 |
| accuracy |  |  | 0.99 | 160 |
| macro avg | 0.99 | 0.93 | 0.96 | 160 |
| weighted avg | 0.99 | 0.99 | 0.99 | 160 |

**Cross-validation**
|  | precision | recall | f1-score | support |
| --- | --- | --- | --- | --- |
| crocodile | 0.98 | 0.96 | 0.97 | 46 |
| cut | 1 | 1 | 1 | 489 |
| accuracy |  |  | 0.99 | 535 |
| macro avg | 0.99 | 0.98 | 0.98 | 535 |
| weighted avg | 0.99 | 0.99 | 0.99 | 535 |

**MAML**
|  | precision | recall | f1-score | support |
| --- | --- | --- | --- | --- |
| crocodile | 0.78 | 0.88 | 0.82 | 8 |
| cut | 0.99 | 0.97 | 0.98 | 73 |
| accuracy |  |  | 0.96 | 81 |
| macro avg | 0.88 | 0.92 | 0.9 | 81 |
| weighted avg | 0.97 | 0.96 | 0.96 | 81 |

**Baseline model on new testing set**
|  | precision | recall | f1-score | support |
| --- | --- | --- | --- | --- |
| crocodile | 0.81 | 0.85 | 0.83 | 20 |
| cut | 0.95 | 0.94 | 0.95 | 65 |
| accuracy |  |  | 0.92 | 85 |
| macro avg | 0.88 | 0.89 | 0.89 | 85 |
| weighted avg | 0.92 | 0.92 | 0.92 | 85 |

In order to test Hypothesis 1 (quality of images impacting the results), the same model applied to the grayscale-augmented dataset produced a similar result to the baseline model: accuracy (98.75% of correct classification of the validation set) remains high, and loss (0.0214) remains low, clearly demonstrating marginal to nil impact of the image quality distribution of the dataset (Table 1). The grayscale-augmented model is more reliable at classifying the highly unbalanced dataset because of the slightly lower loss. Only two crocodile tooth marks were misclassified out of the 160 image validation set (Table 5). This also supports the inference that the blurry and out-of-focus portions of several images may have acted as a regulatory mechanism improving the training of the baseline model (Figure 1, upper). Hypothesis 1 is, thus, rejected.

**Figure 1.**
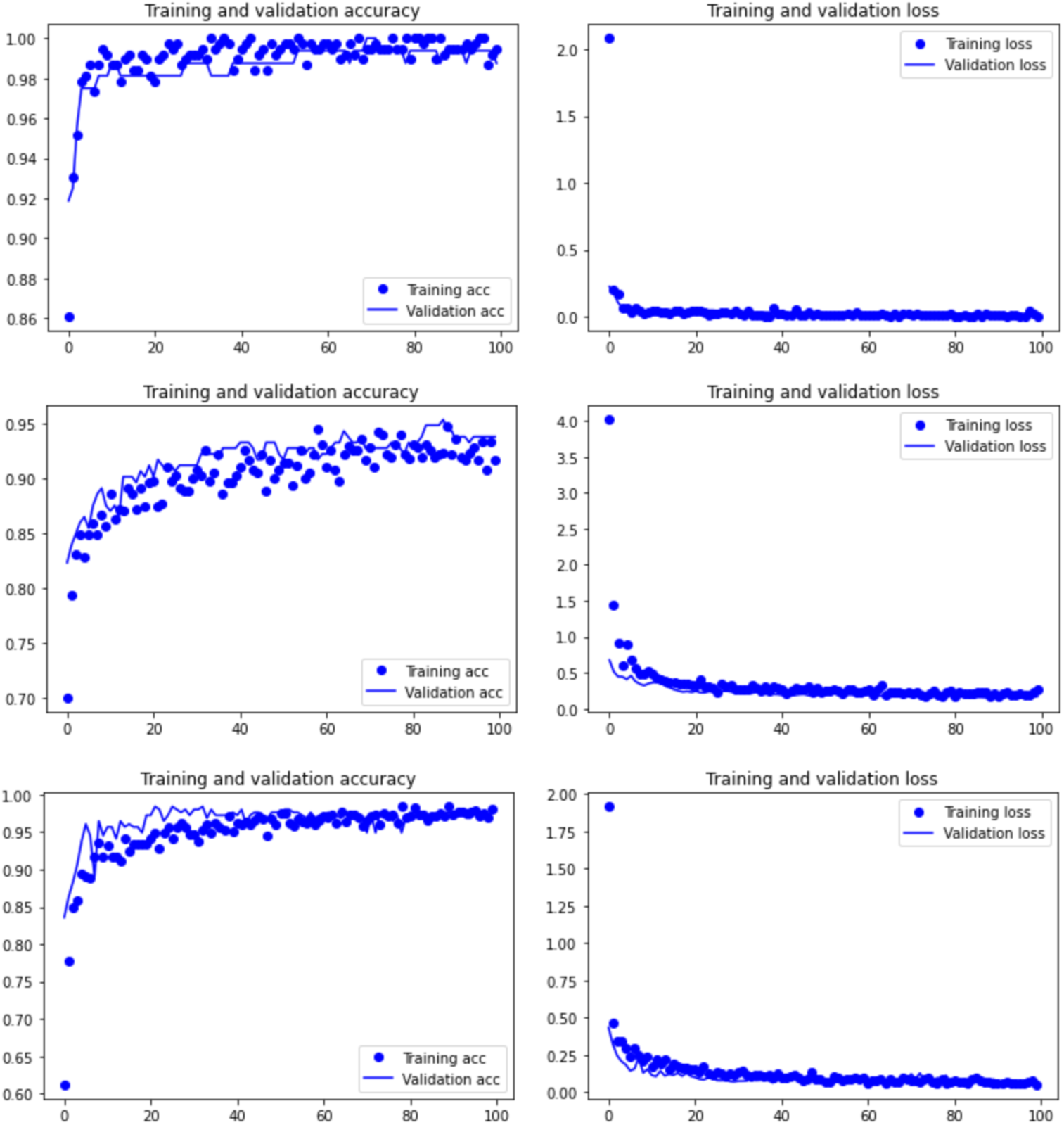
Training graphs-accuracy (left) and loss (right)-of the grayscale intensity-augmented models for the three datasets: Resnet 50 on Dataset 1 (upper), and Dataset 3 (lower), and VGG 16 on Dataset 2 (middle).

The model generated to test Hypothesis 2 (overfitting of the validation set and the resulting model), shows that the four-fold cross-validation resulted in an average accuracy of 99.44%, slightly higher than the baseline model, suggesting lack of overfit. The overall classification report yielded a well-balanced high classification, despite the large difference in sample sizes for both classes (Table 1). The similar results in these metrics (with a substantially larger and more varied validation set) to the grayscale-augmented model indicates that the training-validation method does not show any signs of bias or overfitting. Again, classification of BSM is highly accurate, with only two crocodile tooth marks and one cut mark misclassified (Table 5).

To ensure that cross-validation was not affected by any undetected overfitting process, the MAML analysis, using a separate training-validation-testing subsamples, yielded an accuracy of 96.3% on the testing set, thus supporting that the original model (Abellán et al., 2022) and the baseline model in the present work have not overfit the training or validation datasets (Figure 2). The performance metrics indicate a balanced precision, recall and F-1 score for both classes (Table 1). These metrics together suggest the model generalizes well to unseen data without significant overfitting, and that the model’s predictions are confident for both correct and incorrect classifications. The MAML results are very similar to the cross-validated analyses, further supporting that the train-validation split did not introduce any bias. For this dataset, Hypothesis 2 is also rejected.

**Figure 2.**
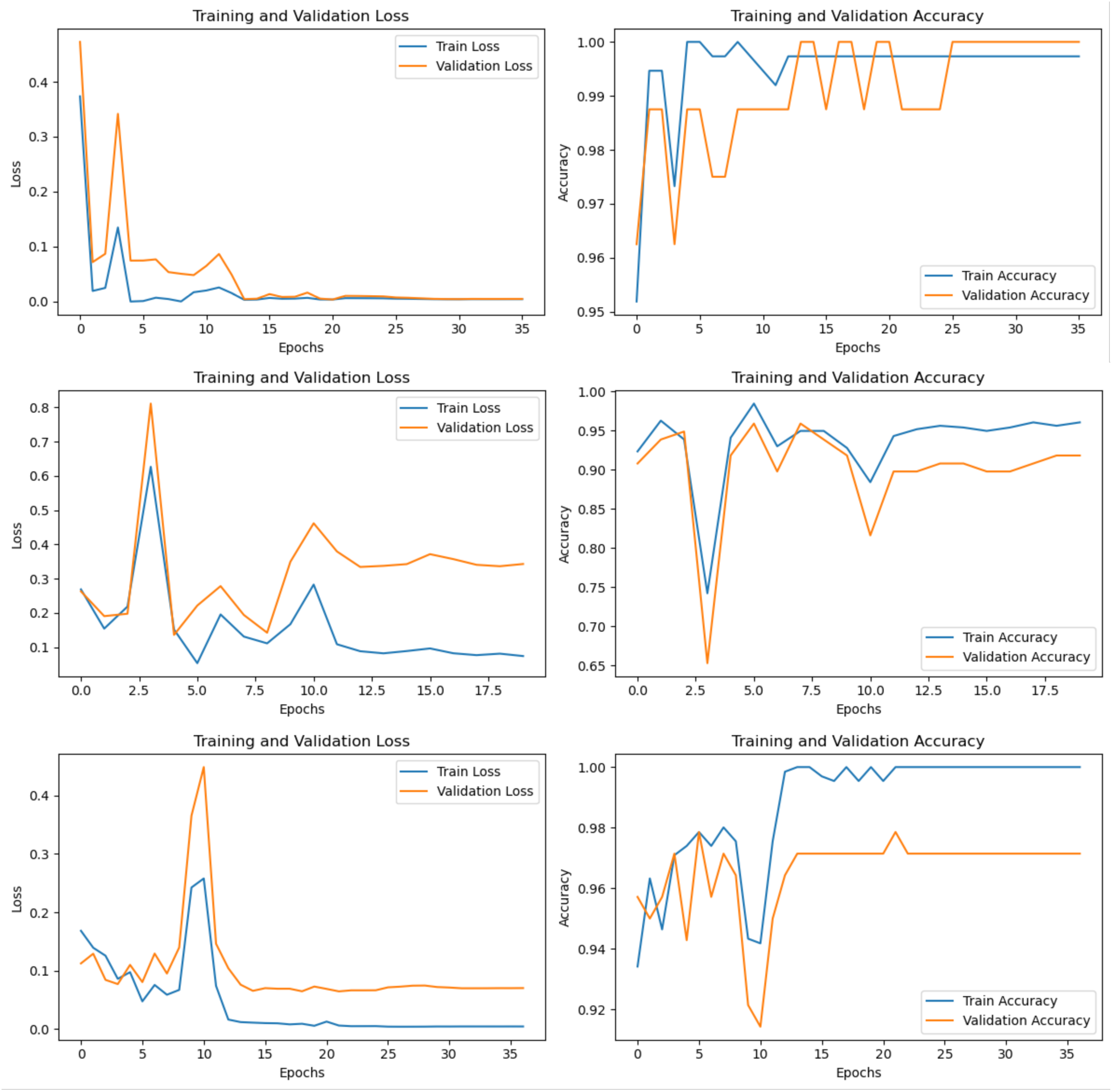
Training graphs of the loss and accuracy of some of the MAML models for Dataset 1 (upper), Dataset 2 (middle) and Dataset 3 (bottom).

Finally, the application of the baseline model to the new testing set composed of 85 new BSM yielded a similar result to all the models described above. A total of 92% of the testing marks were correctly classified, with a F1-score of 0.83 for the crocodile tooth marks and 0.95 for the cut marks (Table 1). Only 3 crocodile tooth marks out of 20 were misclassified, and only 4 out of 65 cut marks were misidentified (Table 7). Interestingly, the three misclassified crocodile marks are the most similar to cut marks of the whole set, by showing extremely narrow V-shaped grooves caused by the carina of the tooth, and resulting from the new crocodiles (*Osteolaemus tetraspis*), which were smaller and with sharper teeth than the original crocodiles (*Crocodylus niloticus)* used for the sample collected from Faunia (Figure 3). These marks would also have been interpreted as cut marks by experienced human taphonomists^3^. This separate testing set shows that the original model learned to differentiate both types of BSM without methodological biases.

**Figure 3.**
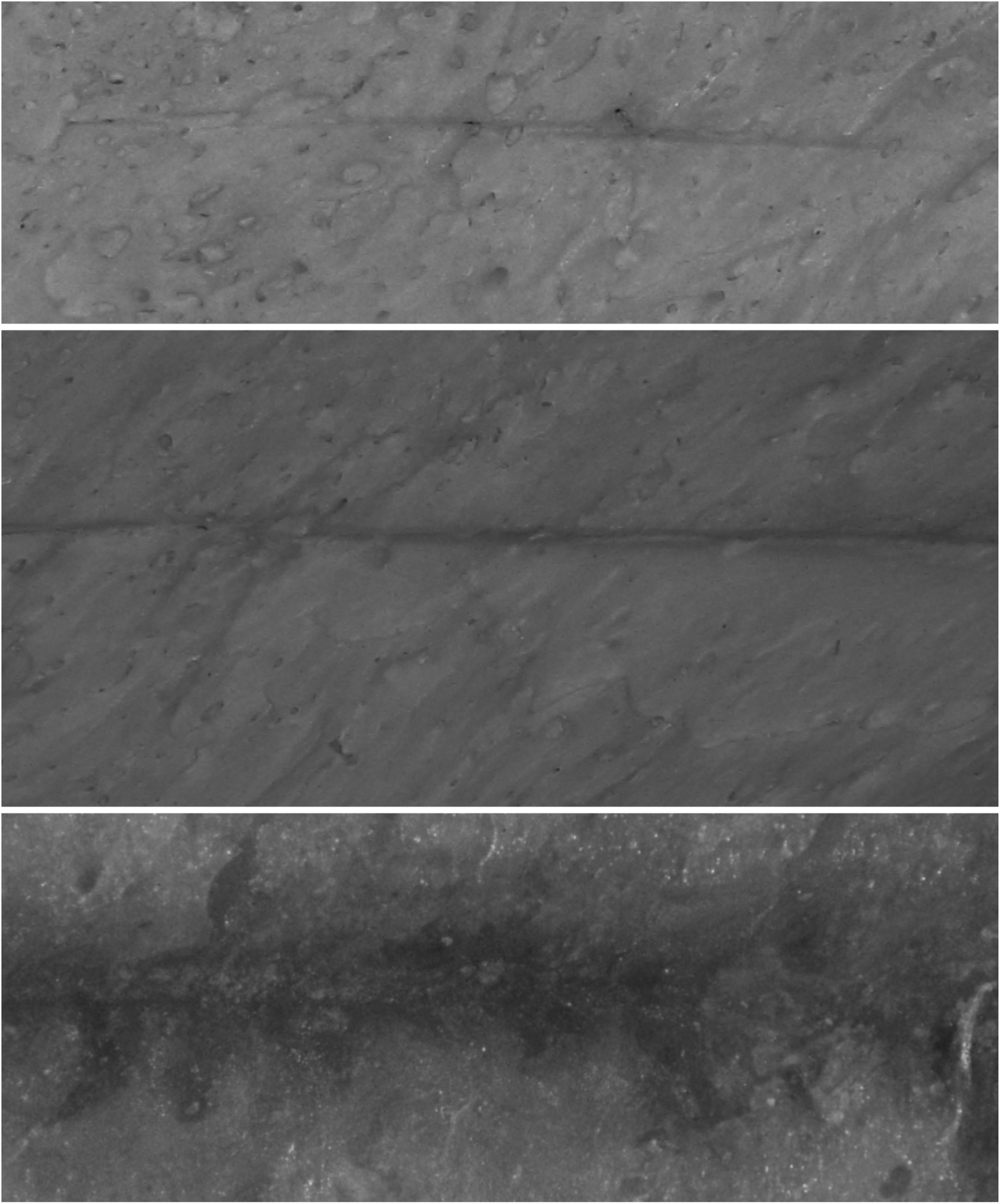
The three misclassified crocodile tooth marks of the new testing set. The two upper marks are from *Osteolaemus tetraspis*, and the bottom one is from *Crocodylus niloticus*. Notice their overall similar morphology to cut marks.

### Dataset 2

A trained VGG16 model yielded an accuracy on the classification of the validation set of 94.27% (loss= 0.140). This is slightly better than the accuracy (0.92) and loss (0.36) scores in the original study (Domínguez-Rodrigo et al., 2020); probably because of the implementation of regularization. High accuracy and moderately low loss together suggest that the modeĺs predictions are fairly confident and close to the true labels, with the exception of trampling.

Initially, this does not suggest overfitting of the model. The performance metrics indicate high precision, recall and F1-scores for all classes but one (Table 2). Tooth and cut marks are fairly well classified (well balanced in their true positives and false negatives). Only 2 tooth marks and 2 cut marks were misclassified in a 193 image validation set (Table 6). Trampling marks exhibit moderate precision (0.79) and recall, with only 61% correctly classified; however, the F1-score of 0.69 indicates a reasonable balance between precision and recall given that subsample size.

**Table 2.**
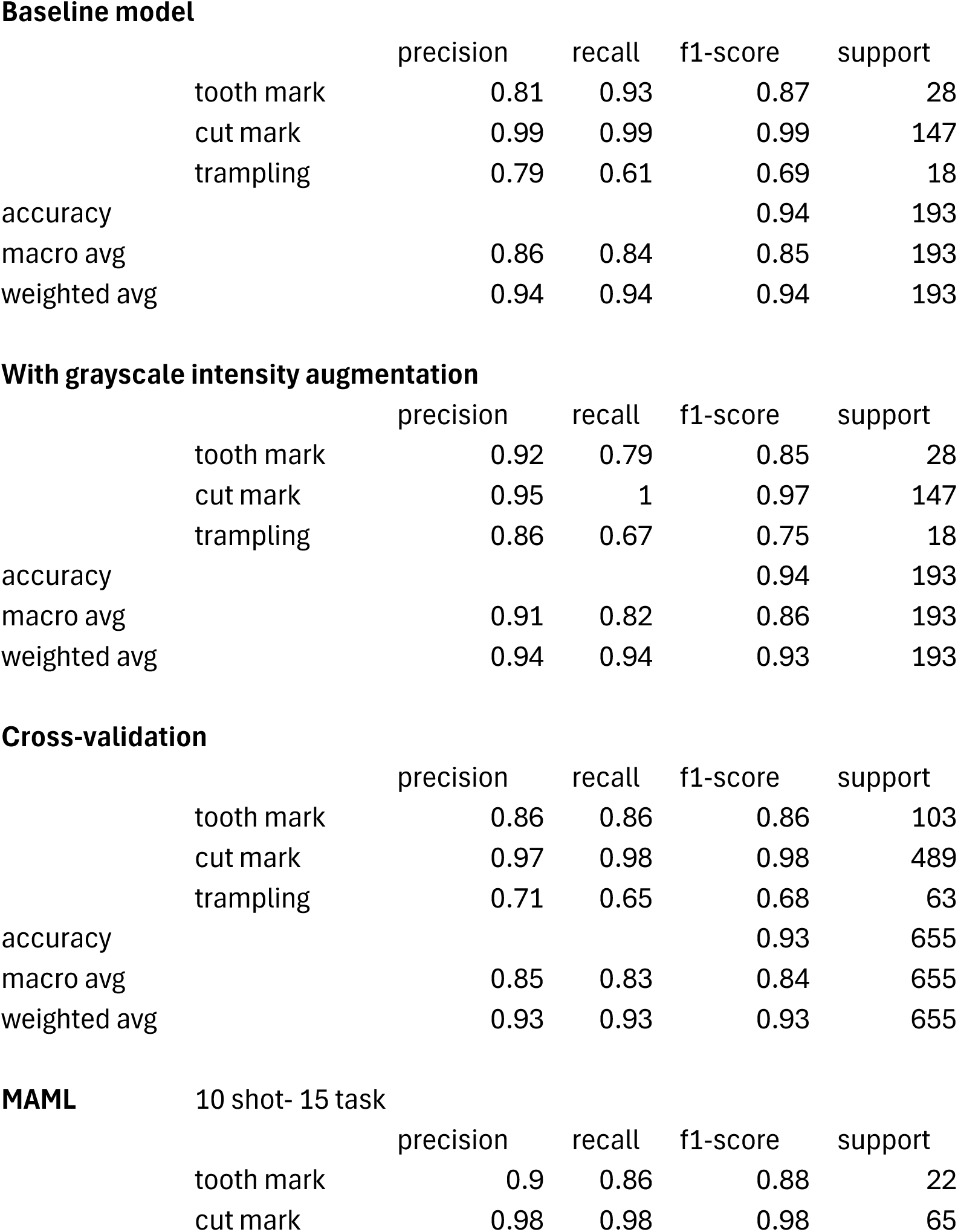
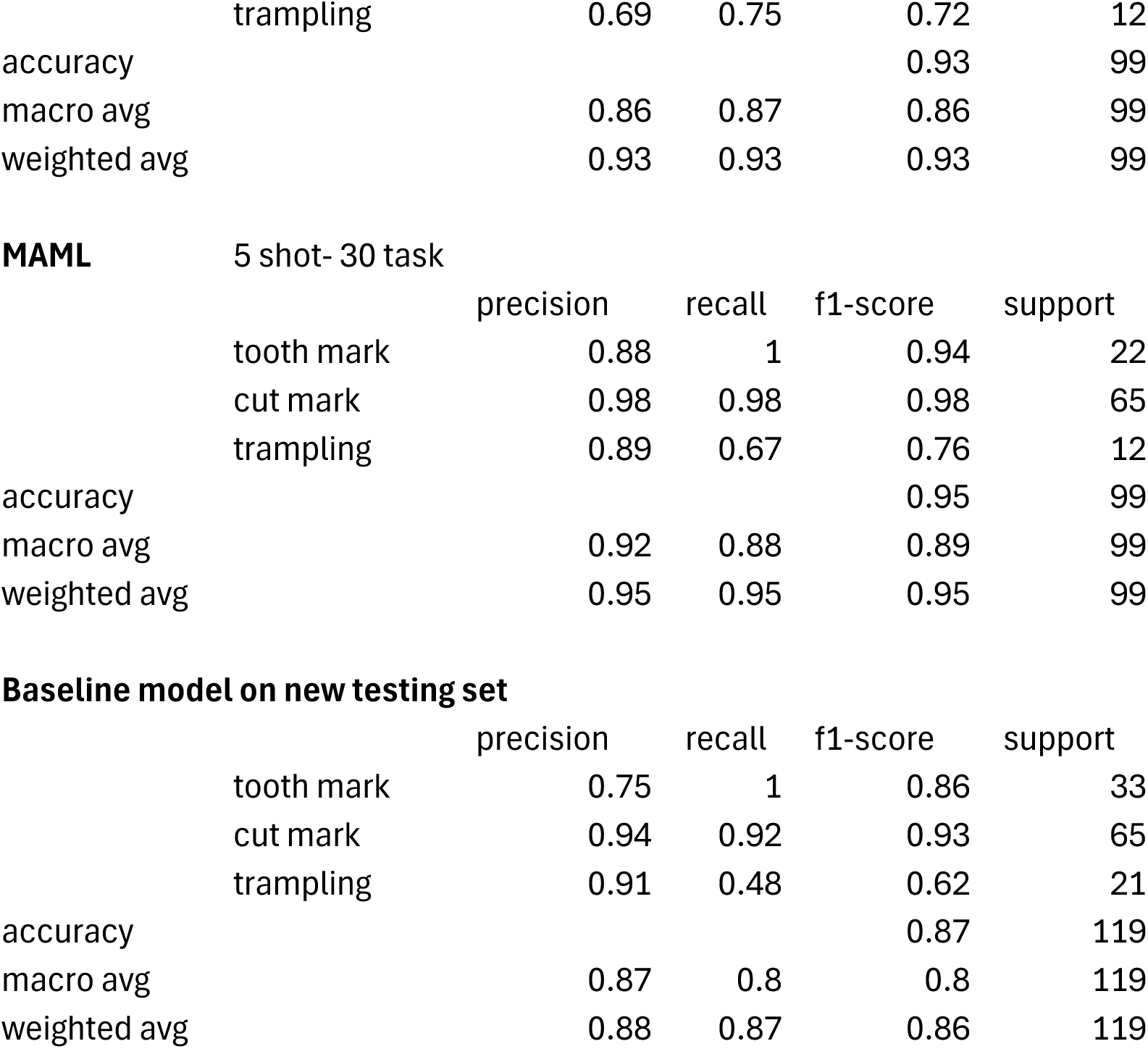
Classification report of the performance metrics from Dataset2 (support=193).

**Baseline model**
|  | precision | recall | f1-score | support |
| --- | --- | --- | --- | --- |
| tooth mark | 0.81 | 0.93 | 0.87 | 28 |
| cut mark | 0.99 | 0.99 | 0.99 | 147 |
| trampling | 0.79 | 0.61 | 0.69 | 18 |
| accuracy |  |  | 0.94 | 193 |
| macro avg | 0.86 | 0.84 | 0.85 | 193 |
| weighted avg | 0.94 | 0.94 | 0.94 | 193 |

**With grayscale intensity augmentation**
|  | precision | recall | f1-score | support |
| --- | --- | --- | --- | --- |
| tooth mark | 0.92 | 0.79 | 0.85 | 28 |
| cut mark | 0.95 | 1 | 0.97 | 147 |
| trampling | 0.86 | 0.67 | 0.75 | 18 |
| accuracy |  |  | 0.94 | 193 |
| macro avg | 0.91 | 0.82 | 0.86 | 193 |
| weighted avg | 0.94 | 0.94 | 0.93 | 193 |

**Cross-validation**
|  | precision | recall | f1-score | support |
| --- | --- | --- | --- | --- |
| tooth mark | 0.86 | 0.86 | 0.86 | 103 |
| cut mark | 0.97 | 0.98 | 0.98 | 489 |
| trampling | 0.71 | 0.65 | 0.68 | 63 |
| accuracy |  |  | 0.93 | 655 |
| macro avg | 0.85 | 0.83 | 0.84 | 655 |
| weighted avg | 0.93 | 0.93 | 0.93 | 655 |

**MAML** 10 shot- 15 task
|  | precision | recall | f1-score | support |
| --- | --- | --- | --- | --- |
| tooth mark | 0.9 | 0.86 | 0.88 | 22 |
| cut mark | 0.98 | 0.98 | 0.98 | 65 |

|  |  |  |  |  |  |
| --- | --- | --- | --- | --- | --- |
|  | trampling | 0.69 | 0.75 | 0.72 | 12 |
| accuracy |  |  |  | 0.93 | 99 |
| macro avg |  | 0.86 | 0.87 | 0.86 | 99 |
| weighted avg |  | 0.93 | 0.93 | 0.93 | 99 |

**MAML** 5 shot- 30 task
|  |  |  |  |  |  |
| --- | --- | --- | --- | --- | --- |
|  |  | precision | recall | f1-score | support |
|  | tooth mark | 0.88 | 1 | 0.94 | 22 |
|  | cut mark | 0.98 | 0.98 | 0.98 | 65 |
|  | trampling | 0.89 | 0.67 | 0.76 | 12 |
| accuracy |  |  |  | 0.95 | 99 |
| macro avg |  | 0.92 | 0.88 | 0.89 | 99 |
| weighted avg |  | 0.95 | 0.95 | 0.95 | 99 |

**Baseline model on new testing set**
|  |  |  |  |  |  |
| --- | --- | --- | --- | --- | --- |
|  |  | precision | recall | f1-score | support |
|  | tooth mark | 0.75 | 1 | 0.86 | 33 |
|  | cut mark | 0.94 | 0.92 | 0.93 | 65 |
|  | trampling | 0.91 | 0.48 | 0.62 | 21 |
| accuracy |  |  |  | 0.87 | 119 |
| macro avg |  | 0.87 | 0.8 | 0.8 | 119 |
| weighted avg |  | 0.88 | 0.87 | 0.86 | 119 |

**Table 3.**
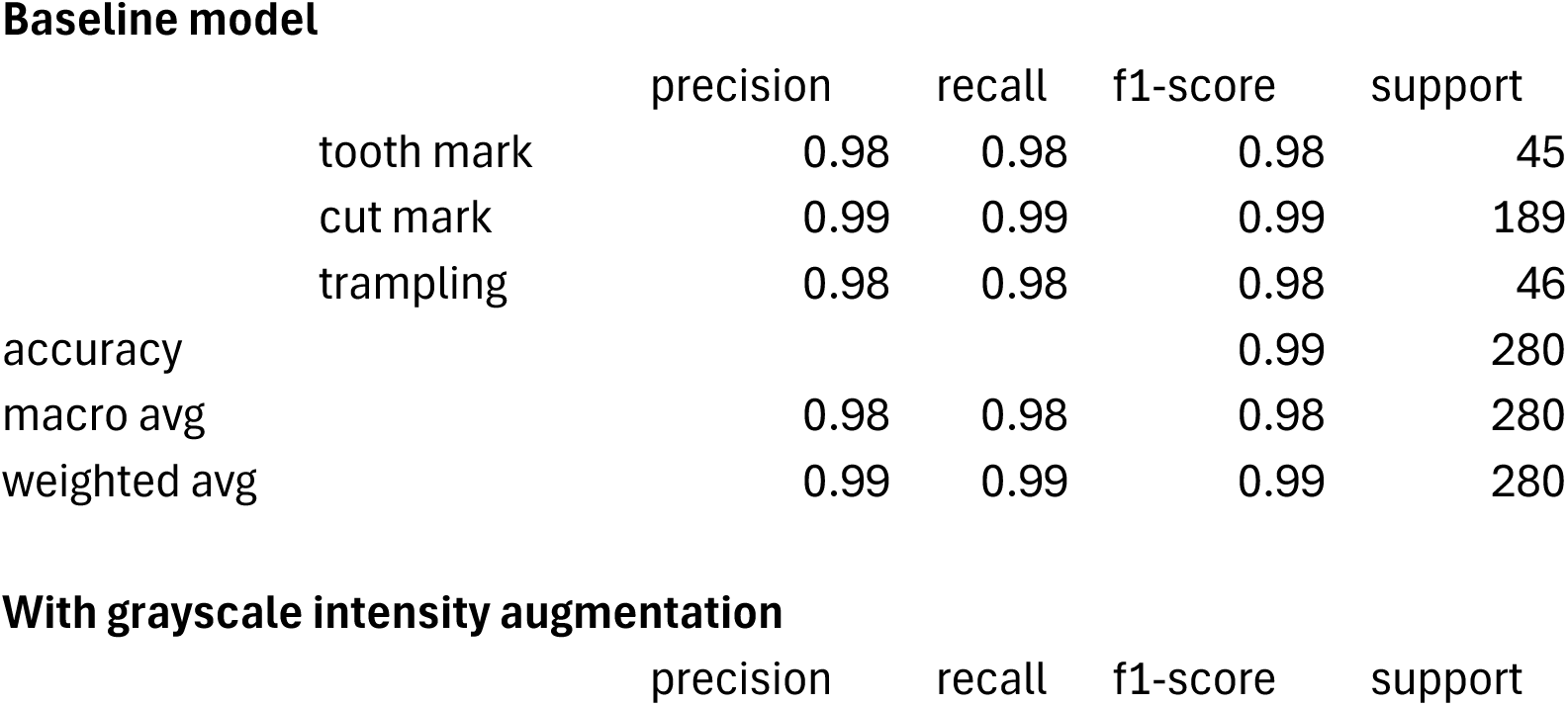
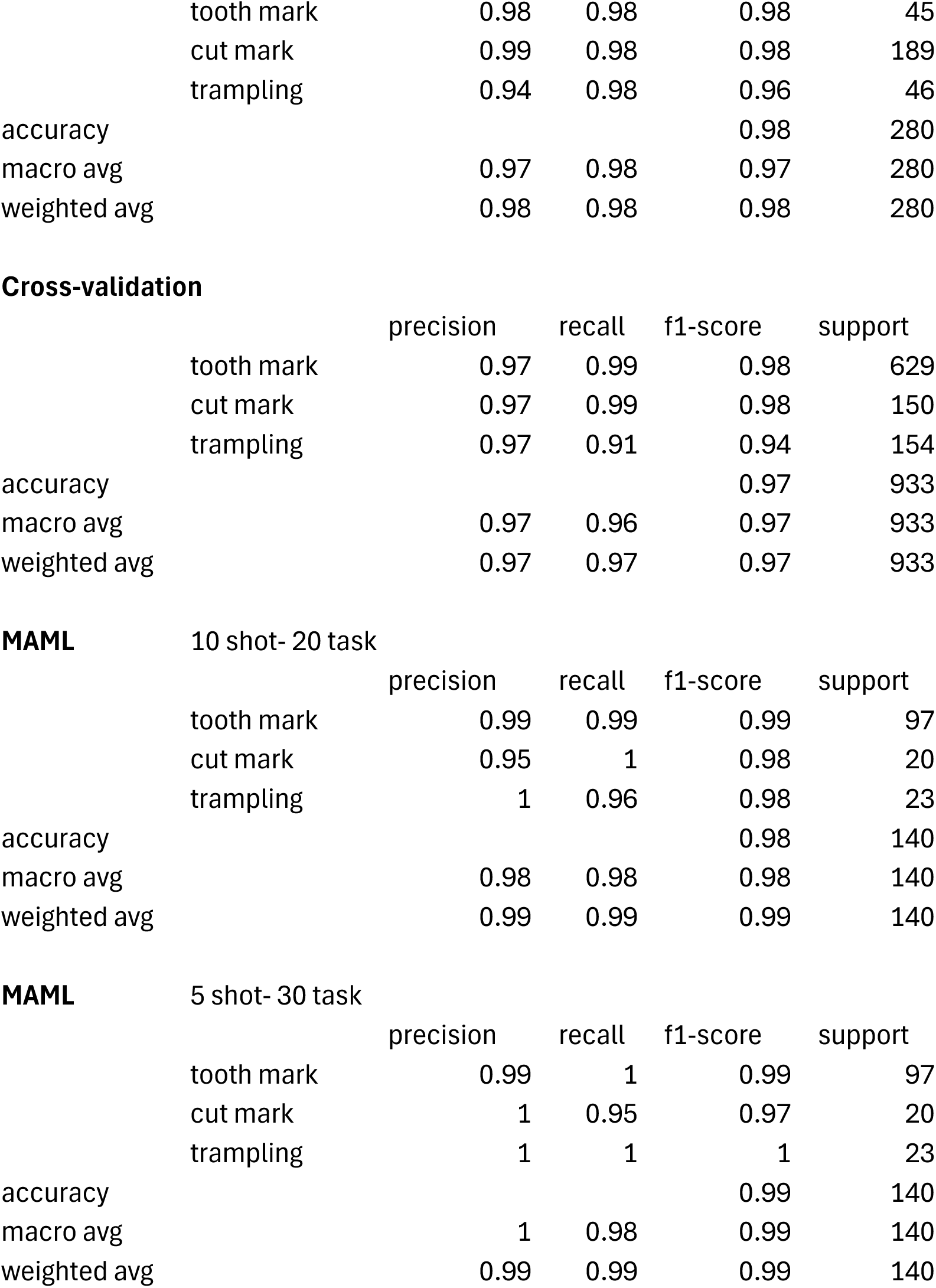
Classification report of the performance metrics from Dataset3 (support=280).

**Baseline model**
|  |  |  |  |  |  |
| --- | --- | --- | --- | --- | --- |
|  |  | precision | recall | f1-score | support |
|  | tooth mark | 0.98 | 0.98 | 0.98 | 45 |
|  | cut mark | 0.99 | 0.99 | 0.99 | 189 |
|  | trampling | 0.98 | 0.98 | 0.98 | 46 |
| accuracy |  |  |  | 0.99 | 280 |
| macro avg |  | 0.98 | 0.98 | 0.98 | 280 |
| weighted avg |  | 0.99 | 0.99 | 0.99 | 280 |
| **With grayscale intensity augmentation** |  |  |  |  |  |
|  |  | precision | recall | f1-score | support |
|  | tooth mark | 0.98 | 0.98 | 0.98 | 45 |
|  | cut mark | 0.99 | 0.98 | 0.98 | 189 |
|  | trampling | 0.94 | 0.98 | 0.96 | 46 |
| accuracy |  |  |  | 0.98 | 280 |
| macro avg |  | 0.97 | 0.98 | 0.97 | 280 |
| weighted avg |  | 0.98 | 0.98 | 0.98 | 280 |
| Cross-validation |  |  |  |  |  |
|  |  | precision | recall | f1-score | support |
|  | tooth mark | 0.97 | 0.99 | 0.98 | 629 |
|  | cut mark | 0.97 | 0.99 | 0.98 | 150 |
|  | trampling | 0.97 | 0.91 | 0.94 | 154 |
| accuracy |  |  |  | 0.97 | 933 |
| macro avg |  | 0.97 | 0.96 | 0.97 | 933 |
| weighted avg |  | 0.97 | 0.97 | 0.97 | 933 |
| MAML 10 shot- 20 task |  |  |  |  |  |
|  |  | precision | recall | f1-score | support |
|  | tooth mark | 0.99 | 0.99 | 0.99 | 97 |
|  | cut mark | 0.95 | 1 | 0.98 | 20 |
|  | trampling | 1 | 0.96 | 0.98 | 23 |
| accuracy |  |  |  | 0.98 | 140 |
| macro avg |  | 0.98 | 0.98 | 0.98 | 140 |
| weighted avg |  | 0.99 | 0.99 | 0.99 | 140 |
| MAML 5 shot- 30 task |  |  |  |  |  |
|  |  | precision | recall | f1-score | support |
|  | tooth mark | 0.99 | 1 | 0.99 | 97 |
|  | cut mark | 1 | 0.95 | 0.97 | 20 |
|  | trampling | 1 | 1 | 1 | 23 |
| accuracy |  |  |  | 0.99 | 140 |
| macro avg |  | 1 | 0.98 | 0.99 | 140 |
| weighted avg |  | 0.99 | 0.99 | 0.99 | 140 |

In order to test Hypothesis 1, the same model was applied to the grayscale intensity-augmented dataset. The result clearly shows that the original non-augmented dataset was not biasing or overfitting the model. The augmented model yielded 93.75% of accuracy with only 0.174 of loss. This is equivalent to the non-grayscale-augmented model, probably because of the random variations of brightness and contrast that were already present in the original dataset. The augmented model is also equally good at classifying the unbalanced dataset, with the exception of the trampling marks (Table 2). Only 6 tooth marks and no cut mark are misclassified in the validation set. Trampling keeps showing a high rate of misclassification (6 out of 18) but it is slightly better than the baseline model (Table 6). This is due to its contrasting small size compared to the tooth mark and cut mark subsamples. Average metrics also show slightly better values than the baseline model (Table 4). This result demonstrates a lack of impact by the quality of the image dataset (Figure 1, middle). Hypothesis 1 is, again, not supported.

**Table 4.** Average values for each model‘s classification metrics showing the scores from Courtenay et al.‘s models for the three datasets, in stark contrast with the values (in bold) from the baseline models replicated in the present work, as well as the color-augmented, cross-validated and MAML models. For more information according to class see Tables 1-3.

| **DS1** |  |  |  |  |  |  |
| --- | --- | --- | --- | --- | --- | --- |
|  | Courtenay et al |  | Present work |  |  | BM-NTS* |
|  | average | baseline | augmented | cross-validated | MAML (best) |  |
| Precision | 0.46 | **0.99** | **0.99** | **0.99** | **0.88** | **0.88** |
| Recall | 0.5 | **0.93** | **0.93** | **0.98** | **0.92** | **0.89** |
| F1 | 0.48 | **0.96** | **0.96** | **0.98** | **0.9** | **0.89** |
| Accuracy | 0.92 | 0.99 | 0.99 | 0.99 | 0.96 | 0.92 |

| **DS2** |  |  |  |  |  |  |
| --- | --- | --- | --- | --- | --- | --- |
|  | Courtenay et al |  | Present work |  |  | BM-NTS* |
|  | average | baseline | augmented | cross-validated | MAML (best) |  |
| Precision | 0.77 | **0.86** | **0.91** | **0.85** | **0.92** | **0.87** |
| Recall | 0.69 | **0.84** | **0.82** | **0.83** | **0.88** | **0.8** |
| F1 | 0.66 | **0.85** | **0.86** | **0.84** | **0.89** | **0.8** |
| Accuracy | 0.86 | 0.94 | 0.94 | 0.93 | 0.95 | 0.87 |

| **DS3** |  |  |  |  |  |
| --- | --- | --- | --- | --- | --- |
|  | Courtenay et al |  | Present work |  |  |
|  | average | baseline | augmented | cross-validated | MAML (best) |
| Precision | 0.88 | **0.98** | **0.97** | **0.97** | **1** |
| Recall | 0.87 | **0.98** | **0.98** | **0.96** | **0.98** |
| F1 | 0.88 | **0.98** | **0.97** | **0.97** | **0.99** |
| Accuracy | 0.91 | 0.99 | 0.98 | 0.97 | 0.99 |

\*Baseline Model on the New Testing Set

The model generated to test Hypothesis 2 shows that the four-fold cross-validation resulted in an average accuracy of 93.13%, which is very similar to the base model (94.27%), suggesting lack of overfit. The overall classification report yielded a well-balanced high classification, despite the unbalanced sample distribution (Table 2). In this case, F-1 scores for the largest subsamples (tooth and cut marks) are 0.86 and 0.98 respectively, which is similar to the base model (0.85 and 0.99 respectively). The small trampling subsample goes from 0.67 in the baseline model, to 0.68 in the cross-validated one (Tables 2 & 6). As was the case for Dataset 1, Dataset 2 seems to be unaffected by overfitting or train-validation split bias.

Just to be further reassured, the high shot version of the MAML model (10 shot-15 task) with the train-validation-test splits generated an overall accuracy on the testing set (which comprised 99 images from three classes) of 92.93% (loss=0.217) (Figure 2).The MAML analysis replicated the cross-validated analysis by showing the same F1-scores for tooth and cut marks, and only a slightly lower value (0.72) for trampling marks. The virtual identity of the data between the cross-validated (train-validation) and MAML (train-validation-test) datasets clearly indicates that there is no detectable overfit of the train-validation method and that the model has learnt efficiently and very similarly at identifying the two most well represented subsamples (Table 2). When applying a low-shot version (5 shot-30 task), these results improve. The overall accuracy is 94.95% (loss= 0.283), with higher F-1 scores for the three types of BSM. Here, trampling shows a F-1 score of 0.76. Again, trampling is misclassified more than tooth and cut marks, whose samples are substantially larger. Given that we are dealing with DL methods, it is not a surprise that the insufficiently represented subsample (trampling) presents these issues. Despite that, as pointed out in the original study (Domínguez-Rodrigo et al., 2020), what the precision/recall metrics indicated was that the potential problem was to misidentify trampling marks with other BSM, but not the other way around.

Lastly, the application of the new testing set composed of 119 new BSM to the baseline model yielded a similar result to all the models described above. A total of 87% of the testing marks were correctly classified, with a F1-score of 0.86 for tooth marks, 0.93 for the cut marks, and 0.62 for trampling marks (Table 2). All tooth marks (n=33) and most cut marks (60 out of 65) were correctly classified (Table 7). This new testing set indicates that there may have been a slight model overfit on the validation test, resulting in poor classification of trampling mark deficient sample, but with very good classification of tooth (recall=1.00) and cut (recall=0.92) marks. The classification of the trampling marks in the new testing set (Table 7) is very similar to that provided by the validation set in the cross-validated model (Table 6). This particular dataset underscores the superiority of the MTL (MAML) method over the DL model in accurately classifying the three types of BSM on their respective testing sets.

### Dataset 3

This is the most extensive and balanced of the three datasets, although it is still unbalanced for cut and trampling marks. One logical prediction would be that it should, by these reasons, produce the best performing models. As a matter of fact, the baseline model (Resnet 50) yielded an accuracy of 98.44% of accuracy (loss=0.044) (Table 3). This is a slight improvement over the original model displayed by Pizarro-Monzo et al. (2023) (accuracy= 97.5%; loss=0.0641). Only one cut mark, 2 tooth marks and one trampling mark had been misclassified (Table 6). The classification metrics yielded extremely high values, with F-1 scores of 0.98 and higher for the three types of BSM.

The same model applied to the grayscale intensity-augmented dataset produced a good result, with only 6 BSM (out of 280) misclassified (Table 6). The new model yielded an accuracy of 97.66% (loss=0.067) and all the classification metrics are >0.94 (Table 3). This is an overwhelming demonstration that the image quality of the dataset did not bias the original baseline model, and that multiple sources of data acquisition do not handicap model performance (Figure 1, lower). Hypothesis 1 is rejected again.

To further ensure that the training-validation method did not result in a model that had overfit on the validation data, the cross-validation analysis yielded 97.32% of accuracy, with F-1 scores for tooth marks (0.98), cut marks (0.98) and trampling marks (0.94) that also showed great classification values for all classes (Table 3). Trampling, usually the most problematic in Dataset2, had high precision (0.97) and high recall (0.91). This result does not support Hypothesis 2.

To ensure that the high performing values of the baseline and cross-validated models were not biased by the training method, a MAML analysis of the same dataset showed 98.57% (loss= 0.037) of accuracy with a 10 shot-20 task approach, and 99.29% of accuracy (loss=0.049) with a 5 shot-40 task model (Table 3) (Figure 2). All F-1 scores are >0.98 for the former and >0.97 for the latter. The replication of all the classification metrics by both MAML models and the baseline, grayscale intensity-augmented and cross-validated DL models (with an even better classification using testing sets in the MAML models) clearly show that the baseline methods for the DL models were methodologically unbiased. The MAML results also underscore the better performance of MAML over traditional DL in the classification of BSM, as was the case for Dataset 2. The better classification of the testing set also shows this compared to the classification of the testing set by the DL model in Courtenay et al.‘s modeling, which showed slightly lower accuracy rates for cut and tooth marks, but substantially more equivocal scores for trampling marks.

## Discussion

### DL is as powerful as the combination of dataset quality and know-how

Courtenay et al (2024) have built a straw man by selecting some of our most unbalanced and smallest datasets-which were initially published as pilot studies to show the potential of the method-to argue that deep learning is currently inefficient for BSM classification. Here, we have shown that such an assertion is inaccurate. All three datasets in our modeling display high values for classification metrics (precision, recall, F-1) for all the classes involved (Tables 1-4), except for the small trampling mark subsample from Dataset 2. This latter datum is related to two factors: small sample size of the trampling mark set, and extremely similar properties of trampling and cut marks. Taphonomists can relate to this, since differentiating between stone tool-imparted cut marks and trampling can at times be an arduous task. When sample size is substantially larger, the DL models can pick better on the microscopic differences and separation of the two types is much better, as demonstrated with the Dataset 3, with the few misclassified trampling marks interpreted as cut marks (Tables 3-4). We have also noticed this effect in other studies, even with larger datasets, where DL is more efficient than any other method at discriminating carnivore agency, except for the smallest subsamples (i.e., crocodiles compared to other carnivorans represented by much larger datasets) (Domínguez-Rodrigo et al., 2024b). We need to stress, though, that the samples of trampling marks used in the analyzed datasets are still too small to be reliable at a more general scale than that represented by the trained models here.

Here, we argue that the methodological decisions made by Courtenay et al. have led them not to replicate the original studies that they set to critique, but to elaborate new models that are substantially less efficient at learning. They did not test if image quality had any effect on our previous models or on their own models for that matter and, however, they created a whole Discussion section full of untested assumptions about it. They simply assumed that some values for contrast, specularity and blurriness (mostly occurring on the periphery of the image/marks as their own GRAD-CAM examples show) invalidated the image set, even if the main features that serve to identify BSM (Domínguez-Rodrigo et al., 2009) can be clearly seen in the central part of most images with the naked eye. If a human taphonomist can perceive those, it is logical to expect that CV methods will also be capable of capturing those features, as the models described in the present work clearly show. This advantage of the machine over the human expert was tested earlier with a simple and small sample for which the best human expert taphonomists reached 62% of accuracy and the machine-with a poorly trained DL model because of sample size-reached >90% of accuracy (Byeon et al., 2019). Some examples of correctly highlighted microscopic features-when using properly trained models-to identify cut, tooth and trampling marks appear on Figures 4-6. See, nevertheless, our criticism of the use of GRAD-CAM in the Appendix.

**Figure 4.**
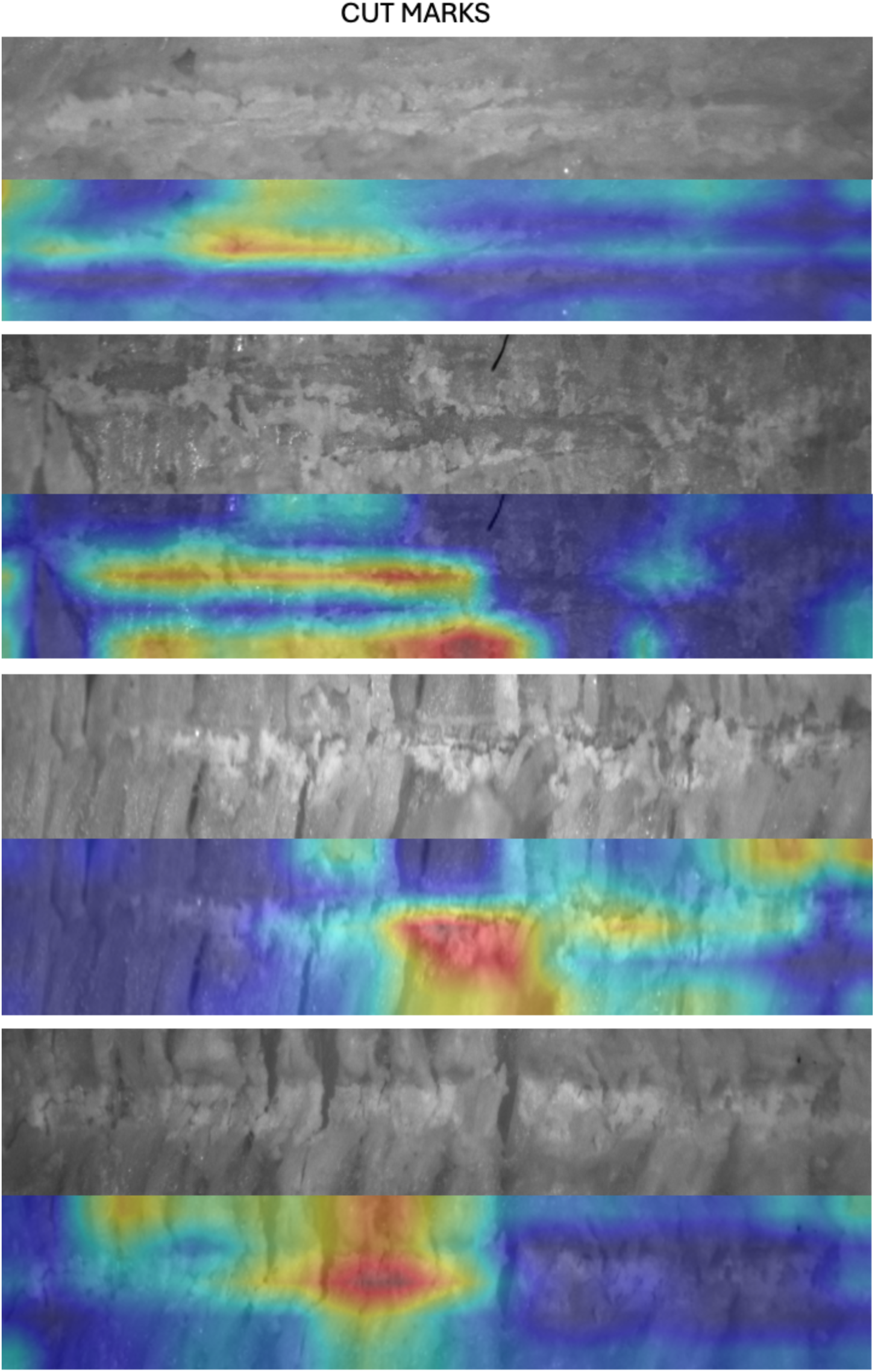
Examples of the GRAD-CAM algorithm capturing relevant parts of cut marks (mostly exfoliation caused by tool crushing cortical bone).

**Figure 5.**
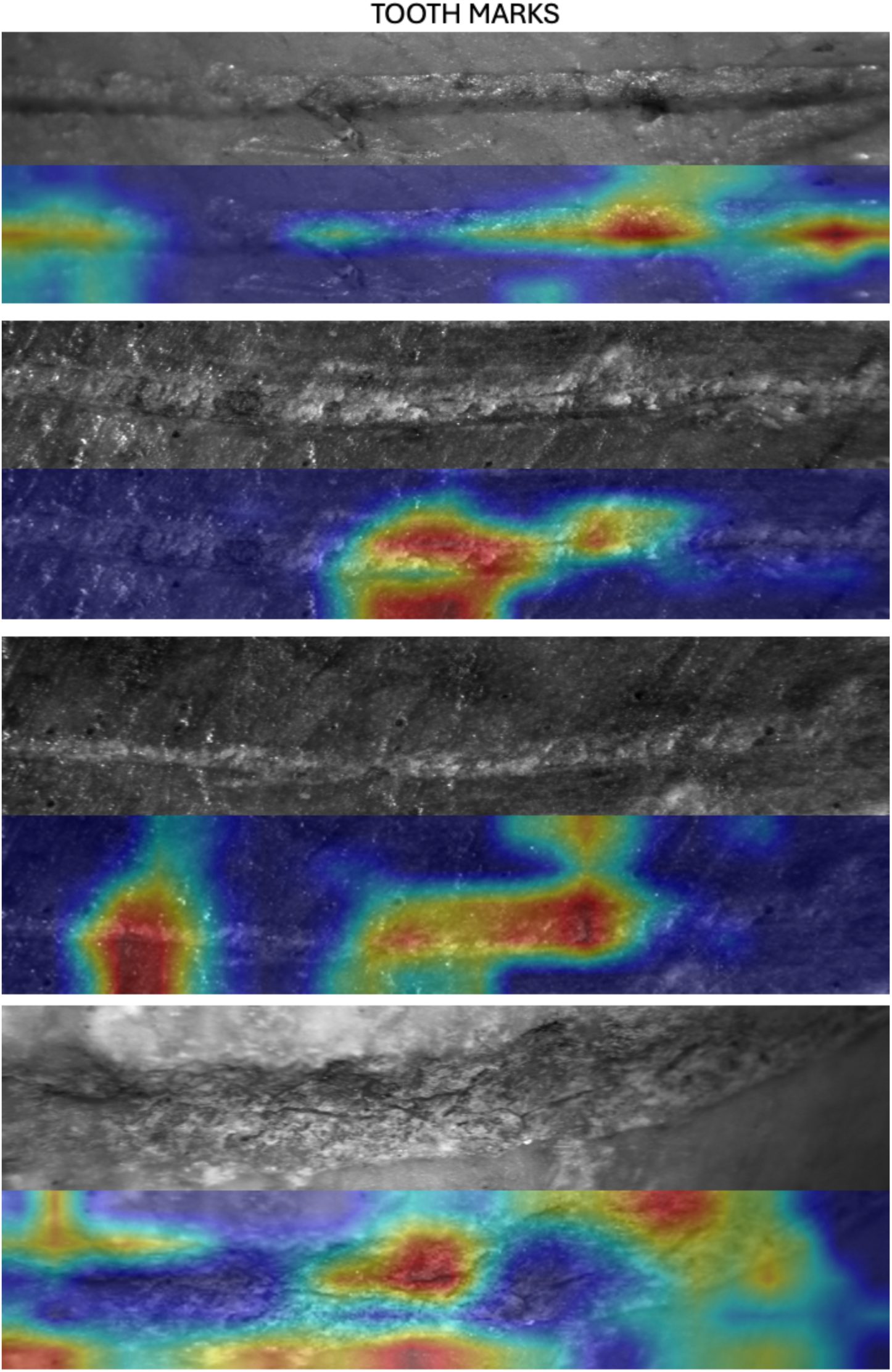
Examples of the GRAD-CAM algorithm capturing relevant parts of tooth marks (mostly crushed cortical bone inside the groove and on its shoulders).

**Figure 6.**
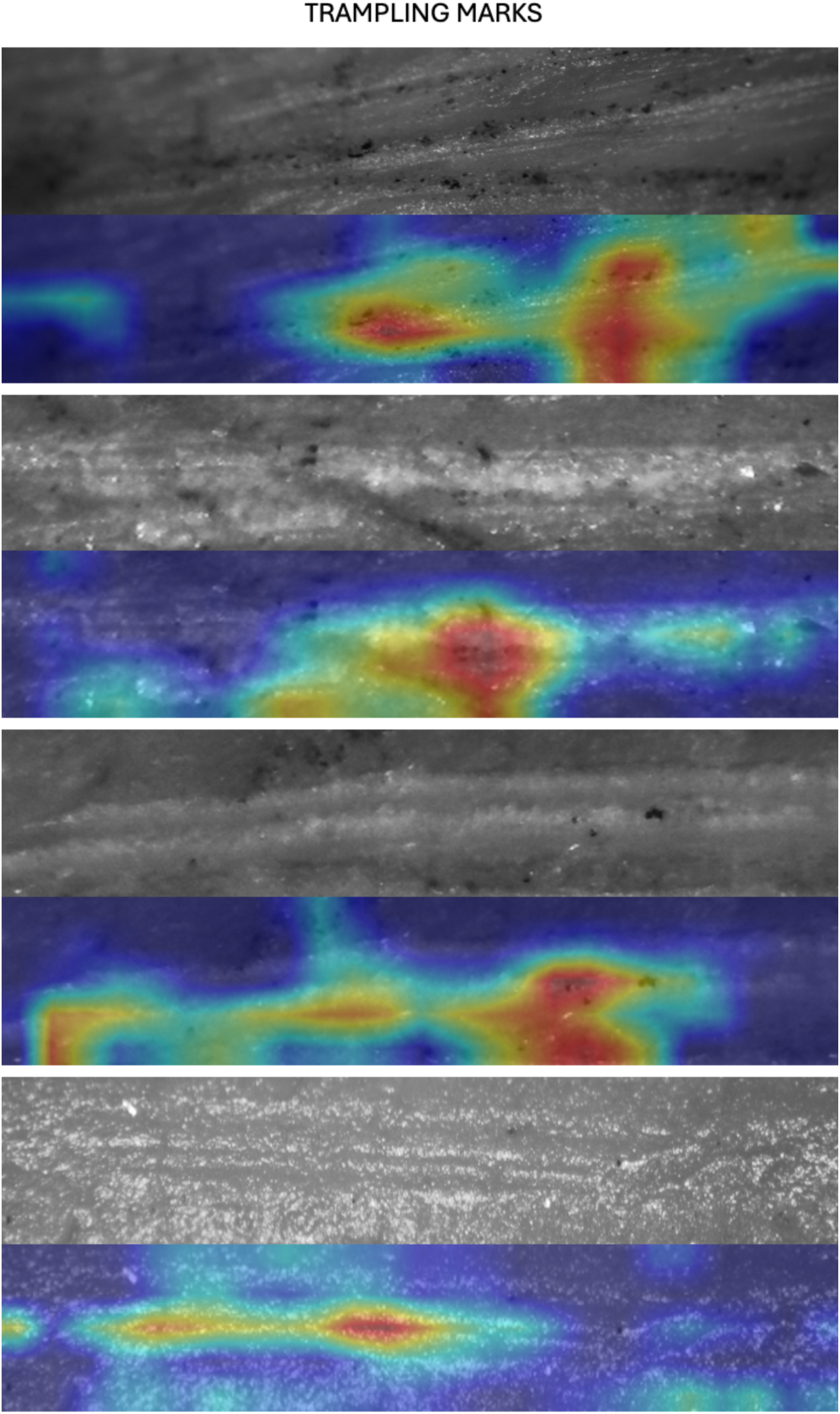
Examples of the GRAD-CAM algorithm capturing relevant parts of trampling marks (mostly the internal part of the groove with microstrations).

Even if Courtenay et al. implemented the same transfer models as the original studies, they reached different accuracy/loss values, because of different methodological choices that they made and different hyperparametrization of those models. These authors followed a traditional triple data split, which is the protocol for representatively large datasets; in this case, they split the three datasets between training (70%) and testing (30%). Training subsets were additionally split, by using 20% for validation. The complete procedure involves: 50% of images assigned to training, 20% to validation, and 30% to testing. This contrasts with the original models of the three datasets, where only training (70%)-validation (30%) sets were deployed. We argue that this decision is in part responsible for the poor performance of their models, because their training and validation sets were substantially smaller, and therefore, less able to learn. Taking into account that this method was applied on extremely small (and unbalanced) samples for DL requirements, the resulting models were probably undertrained/underfit, particularly affecting the classes with the smallest sample sizes. The best example for this could be their modeling of Dataset 1, where they completely misclassify all the crocodile tooth marks (Table 5).

**Table 5.** Confusion matrix for Dataset 1. Notice the sharp contrast between the baseline models in Courtenay et al and the present work. Metrics in bold show the accurate classification values.

| Courtenay et al. |  |  | Baseline |  | Present work<br>Augmented |  | Cross-validated |  |
| --- | --- | --- | --- | --- | --- | --- | --- | --- |
| TRUE | PREDICTED |  | PREDICTED |  | PREDICTED |  | PREDICTED |  |
|  | crocodile | cut mark | crocodile | cut mark | crocodile | cut mark | crocodile | cut mark |
|  | 0 | 13 | **12** | 2 | **12** | 2 | **44** | 2 |
|  | 0 | **146** | 0 | **146** | 0 | **146** | 1 | **488** |

Additionally, training by Courtenay et al. was equally done for all the three databases that they used, even when in the original studies, it is clear that each study used a different set of training parameters. For example, Courtenay et al use mini-batch sizes of 32 for training and validation for the three data sets, when in the original studies these parameters vary, as described in their publications. For the Dataset 1 study, mini-batches were different for training (64) and validation (32) (Abellán et al., 2022). For Dataset 2, both training and validation used mini-batches of size 32 (Domínguez-Rodrigo et al., 2020). For dataset 3, mini-batches also differed for training (32) and validation (20) as explained in the Method section of the original publication, but we noticed that the code included mini-batches of 32 (Pizarro-Monzo, 2023). DL models are highly sensitive to a large array of parameters, and their interplay results in widely divergent results when some are altered. Beyond memory and computational timing, batch size influences the learning process, and it is tightly dependent on the architecture type and the trade-offs between model accuracy and generalization (Hoffer et al., 2017; Hwang et al., 2024; Keskar et al., 2016).

We checked the code that Courtenay et al. used (our access on December 12, 2024 on code that had been last updated on August 26, 2024). There, we realized that another important drawback in their modelś performance was related to the inadequacy of the imaging preprocessing pipeline. There is an unexplainable mismatch between how images have been transformed for training and for validation-testing. For example, for Dataset 2, image pre-processing was made by Courtenay et al using a simple normalization (i.e. 1/255) for the validation/testing sets, and a VGG16 (model-specific) pre-processing function overlaid to simple normalization for the training set. This conflicts with the VGG16 weights used since the original model did not include scaling, and the pixel values of the images in the training set have been transformed differently from those in the validation-testing sets.

Images usually are represented in 8-bit integer format with pixels spanning values from 0 to 255. For better performance, an usual procedure is normalizing the images so that pixel values range between 0 and 1, or-1 and 1 (depending on the models), for faster model convergence. When using transfer learning, models like those used here (e.g., ResNet 50 or VGG16) have pre-processed their image bank in a more complex way than simple normalization. For VGG16, for example, the procedure consists of transforming the pixel values to a normalized range, by the application of a function that subtracts the mean RGB values of the original training dataset (ImageNet) from each pixel in the input image. For each channel, these values are: R (123.68), G (116.779), B (103.939). For Resnet 50, the same mean subtraction is carried out to normalize the input image. This process centers pixel values around zero, which contributes to the stable performance of the gradient descent during training.

The use of inconsistent preprocessing methods for training, validation, and testing datasets—as done by Courtenay et al. —introduces a fundamental mismatch in data distributions across these sets. Specifically, while training incorporates data augmentation and the appropriate preprocessing function of the TL model used, validation and testing images in Courtenay et al.’s VGG16 implementation are merely rescaled (1/255) without applying the same preprocessing pipeline. This inconsistency creates a dual problem: inadequate evaluation of model performance and compromised generalization capability. The inadequate evaluation stems from the distributional mismatch between the training and validation datasets, which differ in pixel value due to the different scaling approaches used caused by the disparate preprocessing steps. This discrepancy leads to an improper assessment of the model’s performance, as the features learned during training may not align with the characteristics of the validation data.

Consequently, validation accuracy and loss metrics fail to reflect the true performance of the model. Moreover, the inconsistency in preprocessing undermines the model’s generalization capability when applied to datasets processed differently, as the trained weights cannot be effectively interpreted by validation or testing data that do not share a homogeneous preprocessing pipeline.

This methodological flaw results in equivocal backpropagation decisions during training, as the model learns features from the augmented training data that are not properly represented or understood by the validation set. Such inconsistencies impair the model’s ability to learn and generalize effectively, as shown in the fact that Courtenay et al. admit that several of their models failed to learn. Pre-trained models, in particular, rely on uniform preprocessing across all data subsets to ensure compatibility with their internal weight interpretations. The failure to maintain preprocessing consistency, as seen in Courtenay et al.’s models, undermines the model’s generalization performance and introduces significant limitations in its practical application. To test the effect of mismatched procedures in the pre-processing of the training and validation/testing subsets, we used different image pre-processing methods in both subsets for the Data Set 1, resulting in all crocodile BSM being misclassified, as shown in Courtenay et al.‘s results.

Similarly, for Dataset 3, Courtenay et al.‘s TL modeling does not include any model-specific preprocessing function in any of the training-validation-testing sets, thus compromising the performance of the model, which was trained using the weights of a model preprocessed with the specific Densenet modeĺs preprocessing function (carrying specific pixel scaling aimed at providing pixel values within the range of [-1, 1], not [0, 1] as performed by Courtenay et al. 2024).

An additional differing issue is introduced by the use of RMSE (Root Mean Square Error) to measure misclassification by Courtenay et al. RMSE is a metric designed to evaluate errors in regression tasks, where the target is the prediction of continuous variables. In classification, with categorical outputs, metrics like accuracy, precision, recall and F-1 are more adequate. RMSE assumes numerical relationships between class labels. This is not relevant, because the distances among classes play no role in classification. Additionally, RMSE cannot be used to understand classification errors, whereas classification metrics can assess how the model is making mistakes through the interplay of true/false positives/negatives. This is the reason why here we used classification metrics and not RMSE.

In summary, the models created by Courtenay et al are methodologically different from those of the original studies. The differences are accounted for: method choice, hyperparameter selection, inconsistencies in image processing for training-validation-testing sets, and different preprocessing functions from those required for TL. It is in the combination of these features that the model pipelines introduce inconsistencies that fatally flaw their results. What these authors succeeded to show is that with their decisions they produced inefficient DL models. For these reasons, their conclusions can only be applied to their own models, and cannot be extended to the original published models nor to those replicated here. This is especially stressed by the fact that in none of our replicated baseline, grayscale intensity-augmented, cross-validated and MAML models did we get accuracy values as low as those reported by Courtenay et al. (2024), and loss values as high, even when using separated testing sets as in our MAML models.

An additional point supporting these interpretations stem from the use of new BSM as external testing sets, which show similar classification metrics to those derived from the baseline training-validation models, the grayscale intensity-augmented models, the cross-validated models and the MAML models. This multiple-source convergence indicates that the DL models function, with trampling marks as the only limiting factor (due to class sample size limitations). Regardless of their differing reliability, the results provided by meta-learning, through the MAML models deployed here, show that in the three datasets, the high degree of accuracy in the classification metrics of all BSM from the testing sets supports that this approach is the most solid for BSM differentiation to date (see next section). This does not mean that the models are good for extensive generalization beyond the characteristics of the BSM modelled. They still serve as pilot studies with a wide margin for improvement, which much bigger image libraries and agencies must exploit for more solid reference models (Cifuentes-Alcobendas, 2025).

We agree that sample size and quality play a major role in sample analysis, but we argue that knowing the limitations of data and the ways to overcome them to maximize information is equally important or more.

### Artificial intelligence turns out to be smarter than we thought

The results that we have presented here show that, regardless of sample size, cut marks and tooth marks are successfully separated by DL models (as documented in the three datasets). Trampling is more problematic in the very small sample size of Dataset 2, but this problem is substantially nuanced in the larger sample of Dataset 3. Dataset 3 is not only the biggest of the three datasets used, but also the one with the best quality images, since those were obtained with much better microscopes (Olympus LEXT OLS3000 confocal microscope, Leica Emspira 3 digital microscope and a KH-8700 3D digital microscope with high intensity LED optics). The fact that the modeling described here yields good discrimination is really encouraging and has been confirmed by Cifuentes-Alcobendas (2025) with an even bigger dataset. The problem with trampling in Dataset 2 was already identified in the original publication: “…trampling marks showing a higher degree of misclassification (recall=30). When misclassified, a greater number of trampling marks are classified as tooth marks instead of cut marks…” (Domínguez-Rodrigo et al., 2020:4)^4^. That model did not learn to identify trampling marks because it was trained on just 63 marks. Despite that, some of the models presented here show that almost 70% of that meagre sample could be correctly classified (cross-validated DL model), and even more, with 75% of correct classification on a separate testing set, if using a MAML model (Table 2). In this particular case, a MAML model is more efficient than a traditional DL model, given that the latter resulted in a model that was unable to classify the new testing set trampling marks as accurately.

It is for this reason that in the past four years an intensive effort has been undertaken in our institution to expand the experimental sample for cut and trampling marks, reaching a total number of 3.272 images of cut marks and 2.412 images of trampling marks. This is by far the largest experimental sample of these BSM available. This has taken 900 hours. Creating an optimized workflow to maximize the performance of CNNs working with BSM image data has required 3,047 hours of computation and 2,356 generated models, resulting from multiple combinations of hyper-parameter tuning and dataset variations. This is also the largest single experiment of parameter combination comparison made for any analysis of DL. It is also the largest dataset for BSM analysis. Given the large sample size, traditional protocols were implemented by creating separate training, validation and testing sets. The latter yielded results of discrimination similar to those shown here for Dataset 3. This is in the process of publication, since the whole process involved a 800-page doctoral dissertation from one of us (GCA) that has just been presented (Cifuentes-Alcobendas, 2025 n.d.). The present work guarantees that DL is substantially up to the task for BSM identification when BSM preservation is good.

In the “Not so intelligent artificial intelligence” section Courtenay et al. engage in a philosophical discussion about the limits of AI, some of which we may agree with, but most of their arguments there are logically flawed. We will stay away from the excessive speculation in that section where very little is empirically supported. All of us working in this field are aware that AI is not human intelligence. This is why we consider inadequate judging the “intelligence” of AI from a human perspective, such as claiming that because “these algorithms are still inherently flawed, lacking fundamental natural cognitive attributes such as common sense” (Courtenay et al., 2024:40), they are imperfect. Let us emphasize that AI is not intelligent in the human sense. AI can be intelligent, much more so, in a different way. AI models outsmart us in many tasks, especially in extremely technical tasks it performs better than humans. AI produces task-specific models that have substantially higher resolution capabilities for pattern finding tasks compared to humans. For example, AI models exceed human radiologists in detecting diseases like breast cancer and lung pathologies in medical scans, offering greater sensitivity and reducing false positives and negatives compared to experts (Ardila et al., 2019; McKinney et al., 2020). They can exceed human ophthalmologists’ diagnostic capabilities for detecting diabetic retinopathy (Gulshan et al., 2016). They also exceed medical experts in detection of skin melanomas (Esteva et al., 2017), brain tumours (Kamnitsas et al., 2017), fracture detection (Rajpurkar et al., 2018), pneumonia (Rajpurkar et al., 2017), cardiac function evaluation (Ouyang et al., 2020), COVID (Wang et al., 2020), cervical cancer (Kurita et al., 2023), degree of osteoporosis (Paderno et al., 2024) by accurately predicting bone density values from CT scan images (Peng et al., 2024), and many other diseases (the literature is too abundant to be summarized here).

All this despite that the quality of x-rays and CT images is heterogeneous and in some datasets even low (Pula et al., 2023; Quaia et al., 2024; Su et al., 2023).

All these AI approaches use imaging and their datasets are derived in many cases from commonly-shared image data banks (because this type of images is so limited) obtained with a myriad of devices, each of them with different digital properties (CT-scan, magnetic resonance, microscopes, medical cameras of a diverse array of manufacturers and with different optical properties). When these images produce models that outperform clinicians, that means that the models work because they thrive on this diverse amount of information. It is for this reason that we approach Courtenay et al.‘s speculation about the effect of digital diversity as distorting and biasing with skepticism. These authors raise the issues of different microscopes having different digital properties that might impact the information of each image and bias DL models. This may be true, but until sufficient experimentation is done, it is sheer speculation. In some of our studies, we opted for combining images from different microscopes to allow models precisely to learn all those nuances (Domínguez-Rodrigo et al., 2024b), just like in AI applications in medical fields. This can probably also justify in part that the models from Dataset 3 produced the more balanced and highest accuracy scores, since their training set includes images obtained by three very different microscopes, with diverse optic properties.

We would emphasize that when using hundreds of BSM images for validation/testing for a task of BSM identification, both DL and MAML models seem to get a much higher percentage right compared to human experts (on much smaller datasets, because human error is increasingly correlated to sample size) who have been doing this for decades (Byeon et al., 2019). In our institution, we have repeatedly done these types of tests. This is not random. It means that the models learn; usually better than us, with all our cognitive attributes, foresight and all our common sense.

### The transferability of CNN learned features

Courtenay et al. argue that large data sets are crucial for AI performance and claim that datasets with 500 images are insufficient because the models/algorithms used contain far more parameters. There are two main objections to this assertion. Whereas the power of a DL model is in general higher when the sample size is very large, that does not mean that DL cannot be efficient with smaller data sets, like the present work has shown. Additionally, this constitutes the biggest contradiction in Courtenay et al.‘s modeling, because they are using small datasets and they are making decisions further reducing the training sets that they use, artificially leading to poorer resolution (see their tinkering of the stacked models in their supplementary files, where they use multiple separate training-validation-testing sets bringing the sample size of each of these subsets to a statistical meaningless point). It seems, in their limited assessment of AI, that Courtenay et al. remain anchored to a view of traditional DL as synonymous with AI, but in the past few years AI has advanced at great length to introduce more modern techniques that are not as data hungry as DL approaches.

Despite their enormous methodological advance (especially in BSM identification, where human experts commonly disagree on the same sets), it is true that the big downside of DL methods is their dependence on big data; i.e., on large amounts of information. Since DL algorithms learn through the use of optimizers that require looping through large batches of data using repeated backpropagation, their performance is tightly linked to the size of the referential database that feeds new information every time. In the past few years, a complementary method called meta-learning (acronym MTL, to differentiate it from Machine Learning [ML]) has been developed to counter the methodological shortcomings when DL is applied to small data sets. MTL basically consists of “learning to learn”. To do so, the MTL algorithms try to mimic human learning. If DL is inspired in the multiple layers of neurons of the human brain, MTL is inspired by the human learning process itself. Humans do not see many images of the same item to properly identify it. They do so with few examples. MTL algorithms are designed with this principle in mind, but using methods that extract information efficiently from a limited set of tasks, which allow subsequently to learn the process of differentiating a new set of tasks. MTL uses algorithms that adapt to these new tasks (even those that they have never seen during training) and have variably good generalization properties. They can outperform DL in generalization when using large datasets, but they do so systematically when datasets are small. The reason is that MTL uses a non-parametric approach, in which training data is dimensionally reduced, memorized and/or embedded into mappings. This contrasts with the parametric approach of other ML methods, which are based on the joint probability distribution of data and their labels.

The most widely used methodological framework for MTL is n-way-k-shot meta-learning (Fei-Fei et al., 2006; Fei et al., 2020; Jadon and Garg, 2020; Li et al., 2017; Ravichandiran, 2018). “Way” translates for “class” and “shot” for “number of supervised examples”. For instance a three-way-five-shot analysis of African carnivores would imply three carnivore species and five examples of individuals within each species. The analysis would involve learning a mathematical representation of each set, with the goal of being able to generalize to other individuals and even other non-trained categories. If we exclude the data augmentation and data generation methods that are also typical of DL, MTL k-shot methods are divided into three basic types: Metric-based, model-based and optimization-based methods (Jadon and Garg, 2020). Metric-based methods are based on mathematical (i.e., metric) similarities between images, which are displayed within an Euclidean space. The key concepts are distances among items and their cosine similarity. The most common architectures are siamese double and triple networks. Siamese neural networks use parallel networks which are trained (by pairing objects and classifying them as similar or dissimilar via a contrastive or a triplet loss function) and tested on two different sets, with the embedding resulting from training applied to the testing features. Matching neural networks are also metric-based. They are constructed using an embedding extractor, whose method relies on LSTM (Long-short-term memory), a cosine similarity distance function and an attention model.

Model-based methods, in contrast, are based on the use of external memory of image features ready to generalize few-shot metalearning. Model-Agnostic Meta-Learning (MAML) is a powerful few-shot learning (FSL) method which is highly adaptable to a wide variety of model architectures. Unlike methods such as siamese networks, prototypical networks, or relation networks, MAML is independent of specific model structures, making it highly flexible for numerous applications (Fei-Fei et al., 2006; Fei et al., 2020; Jadon and Garg, 2020; Li et al., 2017; Ravichandiran, 2018). MAML optimizes model parameters in a way that allows rapid fine-tuning for new tasks, even with minimal additional training data. MAML’s meta-learning approach enables the model to train across multiple tasks, equipping it to generalize quickly to new, unseen tasks. This method contrasts sharply with conventional DL approaches, which typically train on a single dataset (Finn et al., 2017). It is this type of FSL models that we deployed in the present work, with separate training-validation and testing splits. The latter set yielded virtually very similar to almost identical results to those obtained by our training-validation DL analyses. Their classification metrics (precision, recall, F-1) on the three datasets virtually mirror those of the DL models, clearly indicating that there were no inherent biases in the latter by having used training-validation sets. In general, the performance of MAML is more solid, since in the present work, their classification metrics are based directly on a separate testing set that was never used for training/validation. The results of the MAML models alone are enough to certify that this approach is objectively the most efficient at analyzing BSM on bidimensional images.

Our concern about the transferability of DL models is not what Courtenay et al. underline, but the fact that these models have only been trained on exceptionally well-preserved experimental BSM. Their application to prehistoric BSM remains handicapped because most taphonomic BSM have undergone biostratinomic and diagenetic modifications that these models have not been trained to identify (Domínguez-Rodrigo et al., 2025). It is true that in the only two trained models where biostratinomically marks and diagenetically-modified marks, the resolution was high, thereby indicating that there is potential along this experimental route (Pizarro-Monzo et al., 2022; Pizarro-Monzo and Domínguez-Rodrigo, 2020). It is for this reason, that we are switching to experimenting with 3D DL and MTL models, because the true three-dimensional morphology of BSM (not the artificial shapes derived from non-homologous semi-landmarks [see a thorough critique of the limitations of available geometric morphometric studies on BSM in Domínguez-Rodrigo et al., 2025]) may be potentially indicative of agency without having undergone significant transformations during fossil-diagenesis. This still needs to be tested.

We sympathize with Courtenay et al. when they argue that data quality is essential for data interpretation. We also agree that variations in contrast, sharpness, depth of field, in/off-focus may in cases deter models from being efficient. This is the opposite of what has been demonstrated here, probably because most of the low-quality features of the images from Datasets 1 and 2 are the same for all classes (same microscope was used, same illumination, same angle of incidence), and because most of those distortions (with exceptions) occur in the periphery of the cropped images. We argued above that this may have worked as an additional augmentation procedure forcing the models to focus on the most central/clear parts of the images.

Another way in which the efficiency of these DL models is put to test is through confronting agency identification by DL algorithms with that derived from multivariate taphonomic analyses of the same archeofaunal assemblages. For example, a thorough taphonomic analysis of the 1.8 Ma site of DS (Olduvai Gorge, Tanzania) indicated that the agency process in site formation and modification was mostly hominin(primary) and hyenid (secondary) (Domínguez-Rodrigo et al., 2024a). Subsequent DL models applied to the carnivore modifications of the site yielded an overwhelming signal of hyena modifications in the carnivore-modified bones (Cobo-Sánchez et al., 2022). Likewise, a thorough taphonomic analysis of the FLK North assemblages (Olduvai Gorge, Tanzania) indicated a different palimpsestic process in which medium-sized felids had had primary access to carcasses, and hyenids secondary access with marginal to non-existing role of hominins in carcass modification (Domínguez-Rodrigo et al., 2007). The application of DL models trained on different types of carnivores supported this interpretation, identifying the role of leopards and hyenas (Vegara-Riquelme et al., 2023) in the same order and reducing the hominin signal to almost non-existing (Cifuentes-Alcobendas, 2025). The taphonomic analysis of the faunal assemblage from the Upper Pleistocene cave of Tritons (Lleida, Spain) suggested that the accumulation had been made by leopards (Micó et al., 2020). The application of DL models using a modern carnivoran experimental sample led to the identification of the same agency as the primary bone modifier (Jiménez-García et al., 2024). Likewise, the taphonomic analysis of the faunal assemblage from the Toll cave (Barcelona, Spain), indicated that the site was a cave bear hibernation lair with significant activity by carnivores, including scavenging by bears, and potentially other carnivores (Blasco et al., 2020). The application of DL models based on multiple carnivore taxa (including bears) to the BSM sample of the site showed an overwhelming signal of bear modification (with probability >75%) (Pizarro-Monzo et al., 2023). The identification of some cut marks on a hyena phalanx from the Upper Pleistocene Navalmaillo shelter (Madrid, Spain) using traditional multivariate microscopic criteria to separate them from other abrasive agents (Domínguez-Rodrigo et al., 2009) was confirmed by DL models using an experimental dataset including cut marks, trampling marks and tooth marks (Moclán et al., 2024).

If the DL models trained had been as spurious and biased as suggested by Courtenay et al., one would expect a larger variability of contrasting results between the traditional taphonomic analyses of these assemblages and the results provided by the DL studies. The fact that they replicate the same interpretation adds up to their heuristics, and show that the models (despite their incompleteness because of sample size per class and unbalanced nature) did actually learn through their training to identify an important part of the experimental marks, and to transfer their knowledge to efficiently identify unknown but well-preserved prehistoric marks.

We also agree with (Calder et al., 2022) and (Yezzi-Woodley et al., 2024) about the caution in not implementing methods that might lead to data leakage, resulting in biased statistical models. Their criticism of Courtenay et al.‘s (Courtenay et al., 2019) application of Principal Component Analysis (PCA) for dimensionality reduction prior to splitting data into training and testing sets, for example, is correct. They highlight that performing PCA on the entire dataset before splitting can result in data leakage, as the testing set influences the PCA transformation. This contamination skews the evaluation process, leading to artificially inflated accuracy scores and reducing the generalizability of the resulting models to new, unseen data. In contrast, we oppose (Calder et al., 2022) and (Yezzi-Woodley et al., 2024) when they misrepresent the original methods used by other authors. They misrepresented how data augmentation, outlier trimming, and data preprocessing methods were carried out in several works, as exposed by (Moclán and Domínguez-Rodrigo, 2023). They also misrepresented the effects of bootstrapping use on redundant datasets. We are also critical of the way these authors reproduce randomization, which misleadingly inflates the perceived effect of contamination and fails to capture the true patterns in taphonomic data (Moclán and Domínguez-Rodrigo, 2023).

### Equifinality and agency

Courtenay et al. have continuously misused the term “equifinality” in their paper. This is understandably related to these authors not being taphonomists. Only one of them has carried out technical assessment of some archaeological BSM through geometric morphometric analyses of extremely small sample sizes (in all their experimental work, each agent‘s representation is done with samples n=<100 and in some cases as low as n=<30, which falls short of just the morphological diversity of tooth pit types of each of those agents) (Domínguez-Rodrigo et al., 2025). This stays in contradiction with the staunch defense of larger sample sizes made in Courtenay et al. (2024).

Equifinality has two common definitions in taphonomy; one is general, and it is related to two causal processes resulting in *identical* outcomes that are not readily attributable to agent type (Rogers, 2000). This is a taphonomic adoption of the original term, which implied “same final state from different initial states” (Bertalanffy, 1949). The other definition is applied in contexts of two causal processes resulting in *similar* outcomes that are not readily attributable to agent type. The distinction between both concepts is important, because it emphasizes that whereas the first concept is permanent (two identical things cannot be differentiated), the latter is temporary and is contingent on method (Rogers, 2000; Lyman, 2004). It is related to statistical and methodological distinction. Rogers (2000) argues that the use of the term “equifinality” for the second concept is not adequate, because it is not permanent. For example, the issue of agency in faunal accumulations at early archaeological sites was initially interpreted as equifinal, because both hominins and other carnivores could generate similar skeletal profiles.

Later, when fine-grained analysis of skeletal parts was introduced, and quantification of the axial elements could be reliably determined, most of that apparent equifinality vanished (Arriaza and Domínguez-Rodrigo, 2016). In addition, when alternative methods were added (e.g., BSM frequency and anatomical distribution), hominin primary agency in faunal accumulation could be more clearly determined (see summary in Domínguez-Rodrigo et al., 2007; Domínguez-Rodrigo and Pickering, 2017, 2003). We argue that there is room in taphonomy for both definitions, provided no distinction can be grossly made regardless of whether this is a permanent or a temporary state (i.e., open versus closed systems) (Lyman, 2004). Following the first definition, the distinction of BSM, therefore, is not equifinal because no two agents produce “identical” marks, but they can produce similar modifications that only the right method can objectively identify. Instead of equifinality, in those cases there is uncertainty. According to the second definition, BSM are not equifinal because different methods produce different levels of resolution in their differentiation. Some taphonomists using cross-section shape and presence/absence of certain additional features (e.g., microstriations) may not be able to classify marks like others using more complex (e.g. multivariate) methods, involving a much larger array of variables. This would lead to the paradoxical position of BSM being equifinal for some researchers but not to others. The second definition of equifinality does not have to be permanent, but it needs to be general.

The databases used here targeted the differentiation of stone-tool imparted cut marks, carnivore-created tooth marks and sedimentary abrasion in the form of trampling marks. Human taphonomists have for long been able to separate cut/trampling marks from tooth marks. A 30-year-old study on cut, percussion and tooth marks showed that experimental marks of known actor-effector could be correctly diagnosed through blind with accuracy as high as 99% by experts, and novice students could reach 86% of accuracy after three hours of training (Blumenschine et al., 1996/7). Even when using traditional taphonomic microscopic analyses, experimental trampling marks could be identified separately from cut marks 96% of the time.

Therefore, the cut-tooth-trampling BSM differentiation is not subjected to equifinality (the three agents produce morphologically different BSM), but it is the subjectivity of the researcher at capturing those microscopic nuances that make interpretation subjective, with some taphonomists scoring higher than others at detecting true positives (Domínguez-Rodrigo et al., 2017). A more objective method was needed, and this is why AI was brought into taphonomy: to securely identify traces and their agents. The only comparative published study shows that AI does that better than humans, but both are efficient at differentiating BSM, which shows that equifinality (in the sense of identical effects) is not an issue (Byeon et al., 2019).

Only in the case of microscopically close BSM (like tooth marks created by different carnivores, or a few tooth marks created by crocodiles mimicking cut marks) are experts less secure at identifying agency (Abellán et al., 2022; Domínguez-Rodrigo et al., 2024b; Domínguez-Rodrigo and Baquedano, 2018; Jiménez-García et al., 2020; Sahle et al., 2017). This insecurity expands when dealing with prehistoric BSM. This is why the urgency of developing more powerful objective tools to deal with agency (not so much with equifinality) is justified.

## Conclusions

Courtenay et al. (2024) used three previously published small datasets, but failed to replicate the methodologies presented in the original studies, because they adopted methodological decisions that differed from those. These decisions decreased the availability of data to train models, and in the case of their replication of the stacked models in their supplementary files, also a decrease in the availability of data to test models. They assumed that such models are representative of the field, and judged all taphonomic models derived from AI methods by those three studies. In their attempt to explain the divergences of their purported replication, they resorted to poor quality of the image datasets, and models overfitting on the validation sets because of the lack of a separate testing set. In their modeling, the unbalanced nature of the datasets, when trained with smaller samples (leading to biased or underfit models), led to the BSM with the smallest size to destabilize their trained models because they were unable to learn agency through statistically unrepresentative training and validation sets. Mismatched parametrization, and contradictory pre-processing of training/validation-testing sets may have contributed to it. This is why they were unable to classify a single crocodile tooth mark for Dataset 1, misclassify most trampling marks for Dataset 2 and classify correctly only half of the trampling marks for the Dataset 3 (Tables 5-6). Overall, their classification metrics (precision, recall and F-1) remain low in the first two datasets and <10% lower than the models reported here for Dataset 3. In this process, Courtenay et al. have failed to: a) demonstrate the degree to which the quality of the image datasets has impacted the results of the baseline models; b) to demonstrate that the lack of separate testing set has overfit the baseline models, and c) to provide any convincing argument that their replicates in their supplementary information are actually representing the baseline models.

**Table 6.**
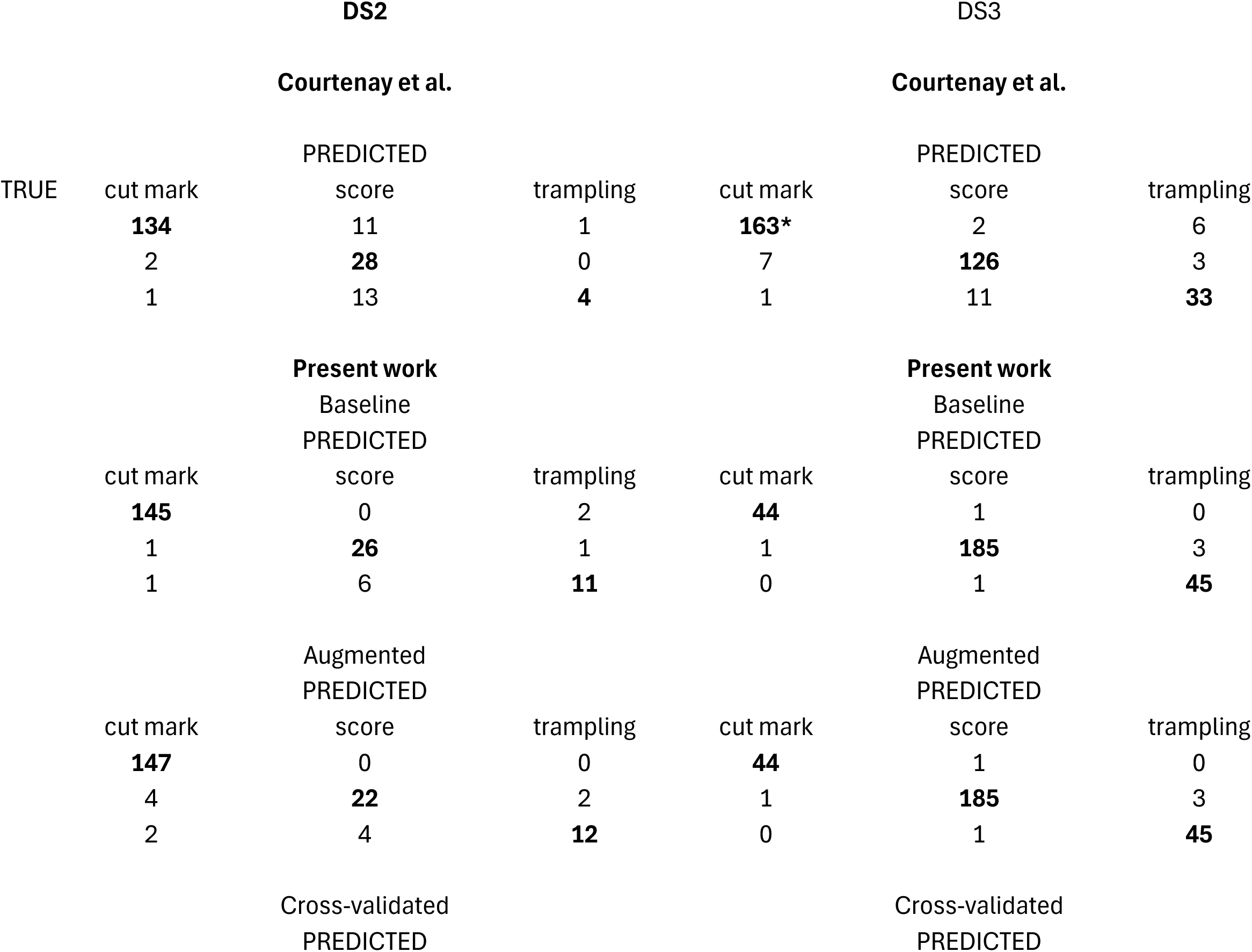
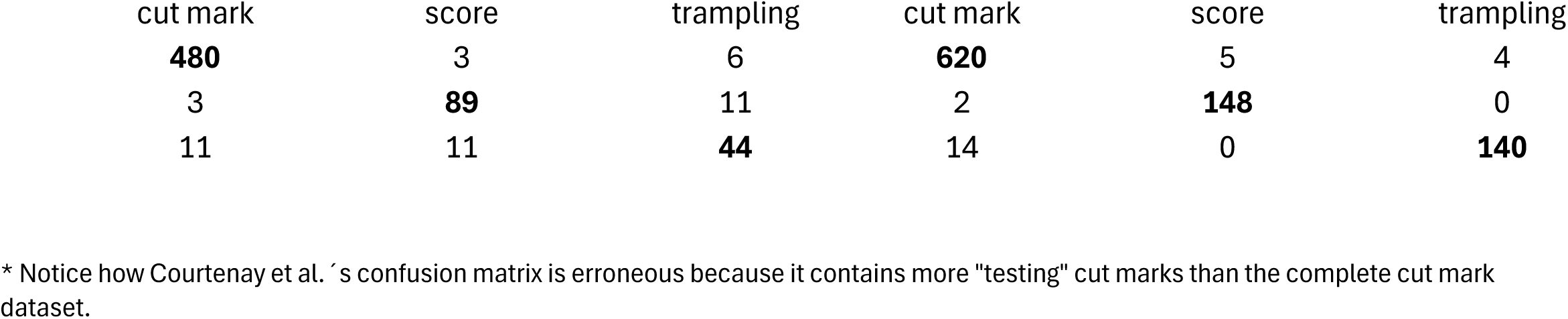
Confusion matrix for Datasets 1 (DS1) and 2 (DS2). Notice the sharp contrast between the baseline models in Courtenay et al and the present work. Metrics in bold show the accurate classification values.

**Table 7.** Classification (confusion matrix) of the datasets using the new testing set.

| TRUE | DS1 |  | DS2 |  |  |
| --- | --- | --- | --- | --- | --- |
|  | BM-NTS* |  | BM-NTS* |  |  |
|  | PREDICTED |  | PREDICTED |  |  |
|  | crocodile | cut mark | score | cut mark | trampling |
|  | **17** | 3 | **33** | 0 | 0 |
|  | 4 | **61** | 4 | **60** | 1 |
|  |  |  | 7 | 4 | **10** |

\*Baseline Model on the New Testing Set

In contrast, we tested the hypothesis that image quality had biased the baseline DL models of the targeted datasets, and rejected it by showing that grayscale intensity-augmented image sets yielded an even higher and more balanced accuracy than the original datasets. The hypothesis that the training-validation method (with the exclusion of a separate testing set) had overfit the models was tested through the most adequate method for small datasets when used in DL modeling: cross-validation. The stratified cross-validated models yielded equal or slightly higher values than grayscale intensity-augmented models and even baseline models (Table 4), showing that the different validation sets used for the training of the k-folds yielded similar accuracy for all classes. New testing sets for Datasets 1 & 2 supported these interpretations. As an additional test to all these models displaying their real learning and generalization potential, a model-agnostic meta-learning (MAML) analysis of the three datasets showed results on independent testing sets similar to or only slightly higher than the baseline, augmented and cross-validated models. This rejects Hypothesis 2, and confirms that the baseline models are unbiased by the factors hypothesized by Courtenay et al. (2024). It also shows that DL and MAML are capable of producing effective BSM models. Even if we accepted at face value Courtenay et al.‘s criticism of previous DL modeling, the results presented here using MAML models clearly show that DL is efficient at classifying BSM. Here, we have shown (as in the original studies) that: a) cut marks and crocodile tooth marks can be reliably differentiated (Abellán et al., 2022), and b) cut and tooth marks can be reliably differentiated, with trampling marks still requiring further work (Domínguez-Rodrigo et al., 2020). This latter interpretation is also supported by Courtenay et al.‘s analysis of Datasets 2 & 3.

These models are functional, but their reliability is limited and their performance is still substantially improvable since their data libraries are small and of poor quality. They should serve as foundation for much more powerful models built upon hundreds of images and more balanced datasets, which are already in the process of being published (Cifuentes-Alcobendas, 2025). Even with these much larger datasets, the generalization capabilities of models will be restricted to the realm of the analogical properties that framed their elaboration. Modellers should be aware that the quality of their experimental sets contains specific substantial, structural and contextual analogical properties that bind them to applications to new data that share these properties (Bunge, 1981). For example, models that have not incorporated biostratinomically and diagenetically modified BSM will not be able to generalize from prehistoric BSM that have undergone morphing through those processes. These samples should contain as much variance as possible to be versatile (instead of restricting it). Only then will they provide the confidence in the identification of BSM in prehistoric and paleontological assemblages that taphonomists are seeking. Criticism of the generalization of the currently available models (within their analogical frameworks), as we have shown here, is largely unwarranted.

## Acknowledgements

We thank the BSM research team at IDEA (Institute of Evolution in Africa), who over years carried out the experiments and introduced the resulting BSM into digital format and the final microscopic analysis. In addition to two authors of this paper (GCA and MVR), these team members are: Blanca Jiménez-García, Natalia Abellán, Marcos Pizarro-Monzo. We are deeply indebted to Edgar Camarós, for letting us use his crocodile taphonomic collection.

## Appendix

### The Limitations of Grad-CAM in Deep Learning Models of BSM

Gradient-weighted Class Activation Mapping (Grad-CAM) is one of several tools in deep learning, used for interpreting convolutional neural networks (CNNs) (Selvaraju et al., 2017). By visualizing the regions of input images that are most influential towards output classification in a model, Grad-CAM enhances our ability to interpret complex models. Despite its utility, Grad-CAM has several limitations that affect its reliability and robustness, because its resolution diverges according to type of task, and the degree of contrasting between delineated features and the remainder of the image. Some of the limitations outlined below are general to any model and dataset, but some are specifically relevant for taphonomic purposes, given that the performance of this saliency method is worst in dull/low-contrast images and in situations of limited class differences. In our experience, the main shortcomings detected in the use of GradCam are the following:

1. *Dependence on Class-Specific Gradients (Predominant-class bias)*. Grad-CAM relies on the diverse gradients of the target class within the last convolutional layer to generate heatmaps. The gradients represent how much the output (e.g., class score) changes when the values in the feature maps (i.e., outputs of the convolutional operations) of the last convolutional layer change. While this approach is effective for clearly defined classes (i.e., morphologically different items), it fails when the model is poorly trained or when dealing with ambiguous or **closely related classes**. In those cases where the distinction between two classes is subtle, the heatmaps commonly fail to highlight meaningful regions, leading to misleading interpretations. For example, in very fine-grained classification tasks, such as differentiating between similar dog breeds, Grad-CAM can fail to highlight meaningful distinctions. If the model struggles to learn subtle differences, the heatmaps may highlight similar regions (e.g., fur or muzzle) for both classes, resulting in ambiguous or uninformative visualizations (Selvaraju et al., 2017). In medical imaging this also happens; it has been observed in tumor classification (Musthafa et al., 2024) or neumo-thorax pathologies (Qiu et al., 2024). There, Grad-CAM can also generate misleading heatmaps when classes overlap significantly. Also, in multi-object images where classes are closely related (e.g., different types of vehicles like cars and trucks), Grad-CAM may generate heatmaps that focus on general features (wheels, body) or even on the spaces between them without clearly differentiating the class-specific features. So, in some of these instances, Grad-CAM is highlighting areas in images that are shared between classes and not class-specific, contradicting its original mission.
2. *Loss of Fine-Grained Localization*. Grad-CAM targets generating coarse visualizations of the regions influencing a model’s decisions. However, its resolution is limited by the spatial dimensions of the last convolutional layer, resulting in heatmaps that may lack fine detail. The image is segmented and highlighted, sometimes containing a diverse array of highlighted elements within the segment. This limitation is particularly problematic for tasks such as object detection and image segmentation, where precise localization of features is critical. Small but important regions in an image, such as fine edges or textures, may be overlooked, reducing the interpretability of the model’s decision-making process. This is extremely important for interpretations of BSM images where the features are not clearly delineated and separated from the rest of the bone (i.e., image), but are an integral part thereof. The lack of fine-grained resolution makes interpretations of Grad-CAM visualizations rather ambiguous and highly subjective when comparing similar classes, since heatmaps in these situations tend to encompass areas larger than those containing the exact defining features (Musthafa et al., 2024). Precise location of complex/ambiguous objects within an image is challenging for Grad-Cam and it may struggle to precisely outline objects due to the coarse nature of the heatmaps.
3. *Sensitivity to Input Perturbations*. One significant drawback of Grad-CAM is its sensitivity to input perturbations. Small changes to the input image, such as noise, variations in lighting, or background clutter, can lead to substantial changes in the generated heatmap. For instance, in medical imaging, slight variations in scan quality could lead to misleading visualizations, potentially affecting critical diagnoses; this, regardless of the resolution of the generated models (Musthafa et al., 2024; Kumar et al., 2023).
4. *Challenges with Multi-Class Problems*. When an image is fed into the model, the output probabilities are calculated for all the classes. In multi-class classification, Grad-CAM focuses on the gradient of the predicted class; that is, the class with the highest probability. This operates by the gradients of the predicted class being computed against the feature map of the last convolutional layer. The gradients then are supposed to reflect the importance of that particular feature map in contributing to the predicted class. Get a different convolutional layer and the gradients will produce a heat map that will activate a completely different part of the image. While this approach works well for visualizing the dominant class, it provides limited insights into other classes. This limitation is particularly problematic when analyzing multi-class images where the goal is to understand how the model perceives all possible classes. In such scenarios, generating separate heatmaps for each class of interest may be necessary to gain a comprehensive understanding of the output, thus incurring again in the aforementioned problems.
5. *Limited Interpretability for Deep Networks*. As networks become deeper, the features captured by the final convolutional layers represent increasingly abstract patterns, which may be difficult to interpret meaningfully, specifically if the classes are morphologically similar. Grad-CAM’s focus on the last convolutional layer means it may ignore important intermediate representations, further limiting its ability to provide comprehensive explanations of a model’s decisions.
6. *Heatmap Resolution Problems*. The spatial resolution of Grad-CAM’s heatmaps is constrained by the size of the feature maps in the last convolutional layer. For models with small feature maps, the resulting heatmaps can be overly coarse, making it challenging to pinpoint relevant regions in the input image. This limitation is especially problematic in high-resolution tasks, where precision is crucial. This is another drawback when using Grad-CAM on images with BSM, where precision of the features targeting identification can become blurry to the method, given their overall integration in the rest of the image (i.e., a mark is only part of the bone, not something that stays in clear contrast against the background) and the coarse nature of the generated heatmaps.

In essence, Grad-CAM is a valuable tool for visualizing and interpreting deep learning models, providing critical insights into model behavior. However, for the purpose of taphonomic applications, its limitations, including dependence on class-specific gradients, coarse heatmap resolution, sensitivity to perturbations, and restricted applicability, highlight its limited value for interpretations of BSM images where:

A. Classes have subtle or overlapping features (point 1), and coarse-grained localization by Grad-CAM fails to detect their boundaries (point 2). This would be the case of a large number of BSM. To a non-expert human, when exposed to the same linear BSM images and asked to point out what they see, most say that they do not see anything, or if they do, they do not delineate the boundaries of marks correctly. Human experts resolve this question better only because they have a mental template of what the marks look like and can identify their boundaries. Trampling and cut marks look so similar that most blind tests with improperly-trained humans are unsuccessful. The degree of resolution necessary to identify the boundaries of similar marks, which occur embedded in cortical bone, is beyond the capability of the algorithm, which is too coarse. For example, in a photo of a savanna with a lion, Grad-CAM can localize the lion but its heatmap will also include other surrounding features that are not part thereof. With BSM, in our observation, we have seen that the heatmap is clearly capturing parts of a mark, but also parts that are unconnected to it, because the boundaries are neither “visible” on the bone surface, nor does the mark stay in contrast with the background (bone surface).

B. Related to the previous point, morphologically similar classes will be differentiated by Grad-CAM with difficulty (point 4). To the naked eye, a tooth score, a cut mark and a trampling mark look very similar or identical, since all of them are linear marks on bone. Expert human taphonomists cannot differentiate many of them without additional magnification. This is a major problem with less distinctive targets like BSM, where the Grad-Cam can sometimes be very inefficient at pointing out marks or even patterned areas. We can find cases where they do so, but most commonly, for all classes Grad-CAM may produce no class-specific pattern.

In some of our previous publications, we have used Grad-CAM, not from our initiative, but upon request by reviewers, despite that we are in general skeptical of their usefulness when dealing with blended images where the targeted object is clearly not separate from its surroundings (i.e., contrasted against the background of the scene).

In the main text, we introduced some images that provide indications that GRAD-CAM is effectively capturing some of the features of the marks. However, the most common pattern is erratic segmentation. The same models that succeed at spotting crucial areas of the BSM to identify class, in other cases fail to do the same and identify completely uninformative segments. For example, Figures 6-8 show how in the cases of cut, tooth and trampling marks, GRAD-CAM is also highlighting segments randomly, which are unaffected by blurring, color contrast, speckles or any other contrasting features.

**Figure 7.**
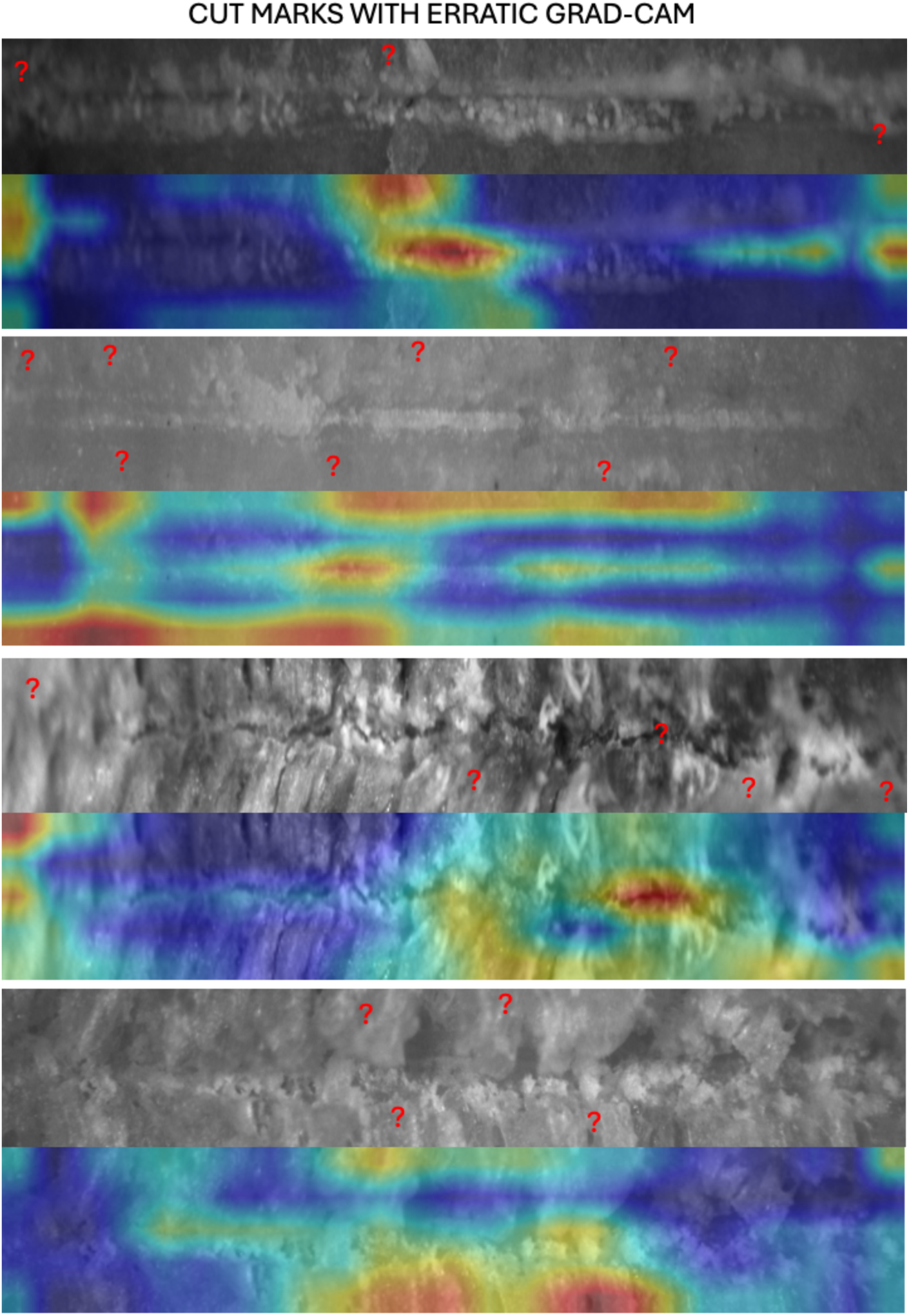
Erratic behavior of GRAD-CAM on cut marks. Notice how the red question marks signal segments that are highlighted without any feature contrasting with the rest of the image. In some cases, the selected area is lighter or darker, but most of the highlighted segments are unaffected by distortion, blurring of unfocused depth of field.

**Figure 8.**
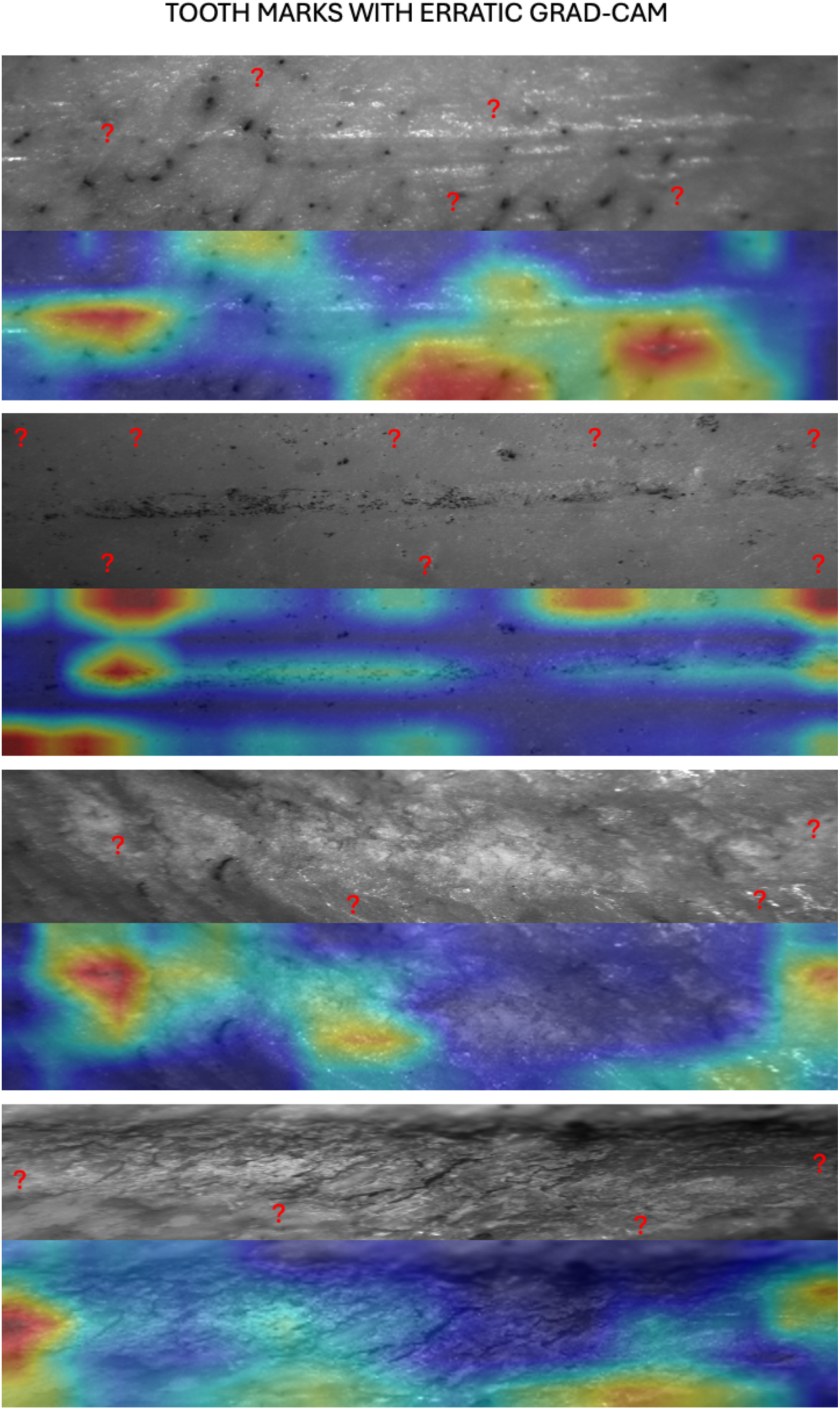
Erratic behavior of GRAD-CAM on tooth marks. Notice how the red question marks signal segments that are highlighted without any feature contrasting with the rest of the image. In some cases, the selected area is lighter or darker, but most of the highlighted segments are unaffected by distortion, blurring of unfocused depth of field.

**Figure 9.**
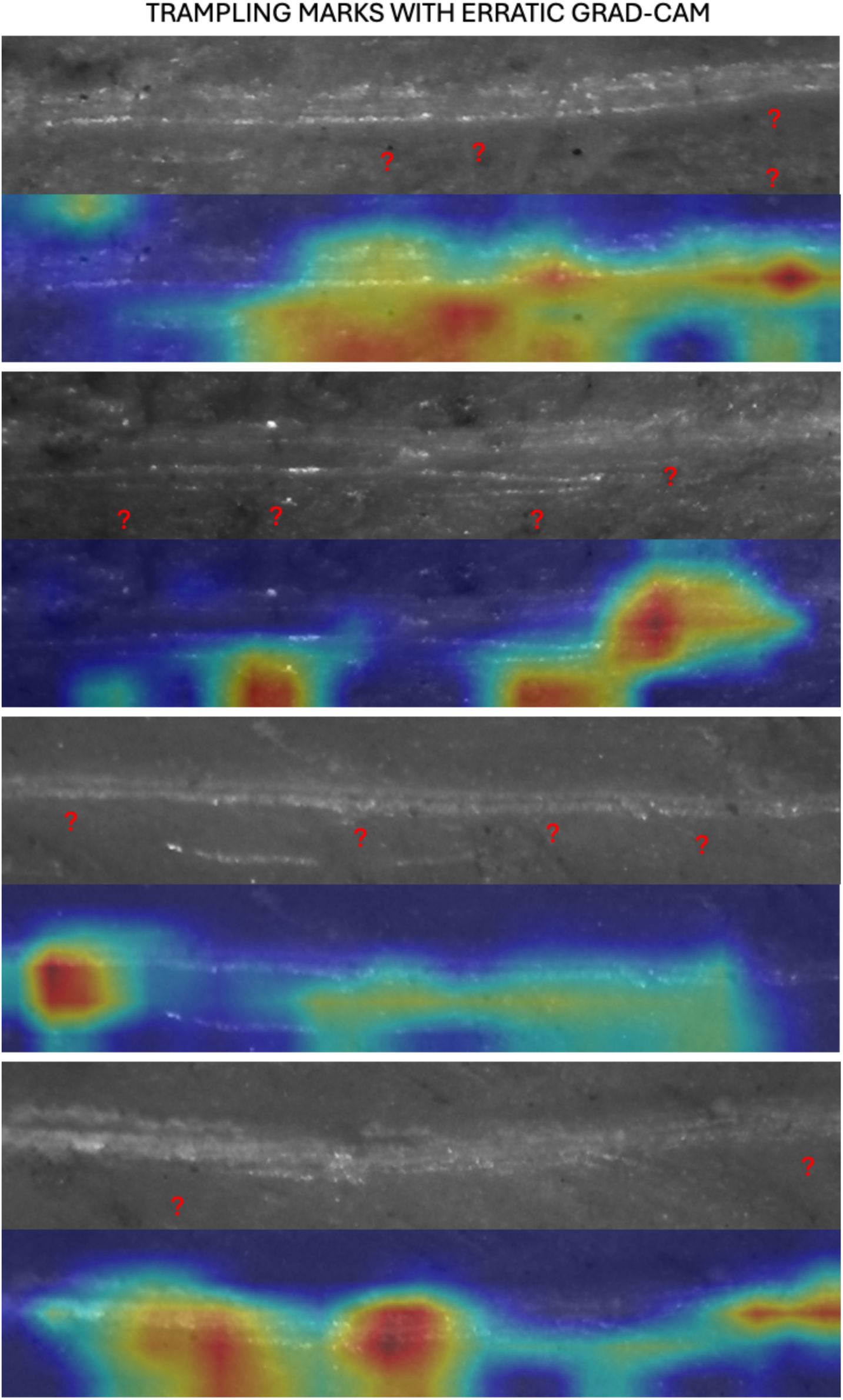
Erratic behavior of GRAD-CAM on trampling marks. Notice how the red question marks signal segments that are highlighted without any feature contrasting with the rest of the image. In some cases, the selected area is lighter or darker, but most of the highlighted segments are unaffected by distortion, blurring of unfocused depth of field.

## Footnotes

1 Notably, several of the studies in which these DL models were originally applied are the same ones featured in publications co-authored by Courtenay (e.g., Cobo-Sánchez et al., 2022; Domínguez-Rodrigo et al., 2022), where no critical remarks regarding their implementation or methodology were previously expressed by this author.

2 “If little data is available, then your validation and test sets may contain too few samples to be statistically representative of the data at hand….K-fold and iterated k-fold validation are two ways to address this” (Chollet, 2022).

3 As a matter of fact, after the completion of this study MDR remarked that he had never seen a crocodile tooth mark with the morphologies shown in Figures 3ab. He remarked that he thought those marks were cutmarks imparted with metal knives during the butchery or preparation of the specimens for experimentation. Their location on the bones where they are documented would support that. If so, this would further reinforce the efficacy of the DL models to discriminate cut marks from crocodile tooth marks, and the accuracy values reported here for the new testing set here would be an underestimation.

4 This was a misinterpretation in the original publication (Domínguez-Rodrigo et al., 2020). Most of the misclassified trampling marks (n=12) were erroneously classified as cut marks and not tooth marks (n=4).

## Notes

https://doi.org/10.7910/DVN/WUSGSW

### Competing Interest Statement

The authors have declared no competing interest.

